# The Cohesin ATPase cycle is mediated by specific conformational dynamics and interface plasticity of SMC1A and SMC3 ATPase domains

**DOI:** 10.1101/2022.06.24.497451

**Authors:** Marina Vitoria Gomes, Pauline Landwerlin, Marie-Laure Diebold-Durand, Tajith B. Shaik, Alexandre Durand, Edouard Troesch, Chantal Weber, Karl Brillet, Marianne Lemée, Christophe Decroos, Ludivine Dulac, Pierre Antony, Erwan Watrin, Eric Ennifar, Christelle Golzio, Christophe Romier

**Affiliations:** Université de Strasbourg, IGBMC UMR 7104 - UMR-S 1258, F-67400 Illkirch, France; CNRS, UMR 7104, F-67400 Illkirch, France; Inserm, UMR-S 1258, F-67400 Illkirch, France; IGBMC, Institut de Génétique et de Biologie Moléculaire et Cellulaire, Department of Integrated Structural Biology, F-67400 Illkirch, France; IGBMC, Institut de Génétique et de Biologie Moléculaire et Cellulaire, Department of Translational Medicine and Neurogenetics, F-67400 Illkirch, France; Architecture et Réactivité de l’ARN, Institut de biologie moléculaire et cellulaire (IBMC), UPR 9002 du CNRS, Université de Strasbourg, 15 Rue René Descartes, 67084 Strasbourg Cedex, France; CNRS, Université de Rennes, Institut de Génétique et Développement de Rennes, UMR 6290, Rennes, France

## Abstract

Cohesin is key to eukaryotic genome organization and acts throughout the cell cycle in an ATP-dependent manner. The molecular mechanisms underlying the Cohesin ATPase activity are poorly understood. Here, we have characterized distinct steps of the human Cohesin ATPase cycle and show that the SMC1A and SMC3 ATPase domains undergo specific but concerted structural rearrangements along this cycle. Specifically, while the proximal coiled coil of the SMC1A ATPase domain remains conformationally stable, that of SMC3 displays an intrinsic flexibility. The ATP-dependent formation of the heterodimeric SMC1A/SMC3 ATPase module (engaged state) favours this flexibility, while it is counteracted by binding of NIPBL and DNA (clamped state). Opening of the SMC3/RAD21 interface (open-engaged state) leads to a stiffening of the SMC3 proximal coiled coil that constricts, together with that of SMC1A, the DNA binding chamber of the ATPase module. Our results reveal that the plasticity of the ATP-dependent interface between the SMC1A and SMC3 ATPase domains enables the structural rearrangements occurring between the engaged, clamped and open-engaged states, while keeping the ATP gate shut.

## Introduction

Structural Maintenance of Chromosomes (SMC) complexes play key roles in genome organization in the three domains of life. While these complexes share a similar architecture and a common ATPase activity, they have evolved different functions supported by specific sets of core, auxiliary and regulatory subunits (Davidson and Peters, 2021; Haering and Gruber, 2016; Yatskevich et al., 2019). How shared and evolutionary-divergent mechanisms of SMC complexes contribute to their specific functions remains poorly understood. Among the SMC complexes, eukaryotic Cohesin displays broad functional implications. Cohesin is notably involved throughout the cell cycle in sister chromatid cohesion, chromosome segregation, DNA repair, chromatin loop formation and, in vertebrates, in V(D)J recombination and in higher order chromatin organization by forming Topologically Associated Domains (TADs). To perform these functions, Cohesin acts in an ATP-dependent manner by synergistically alternating between different mechanisms, including dynamic loop extrusion and DNA tethering (Davidson and Peters, 2021; Yatskevich et al., 2019).

The core Cohesin complex is composed of two Smc subunits, SMC1A and SMC3, and of the RAD21^Scc1^ kleisin subunit (human names used throughout; *S. cerevisiae* names are given in superscripts when diverging significantly from the human names). The SMC1A and SMC3 subunits harbor at each of their extremities a hinge domain and an ABC-Transporter-like ATPase head domain (HD), both separated by a long intramolecular antiparallel coiled coil (Gligoris and Lowe, 2016; Hopfner and Tainer, 2003; Yatskevich et al., 2019). Formation of the Cohesin core complex involves SMC1A and SMC3 heterodimerization through their hinge domains and the asymmetric bridging by the RAD21^Scc1^ kleisin subunit of the SMC1A and SMC3 HDs to form the tripartite ring characteristic of SMC complexes (Figure 1A) (Gligoris et al., 2014; Haering et al., 2002; Haering et al., 2004; Kurze et al., 2011; Shi et al., 2020).

**Figure 1.**
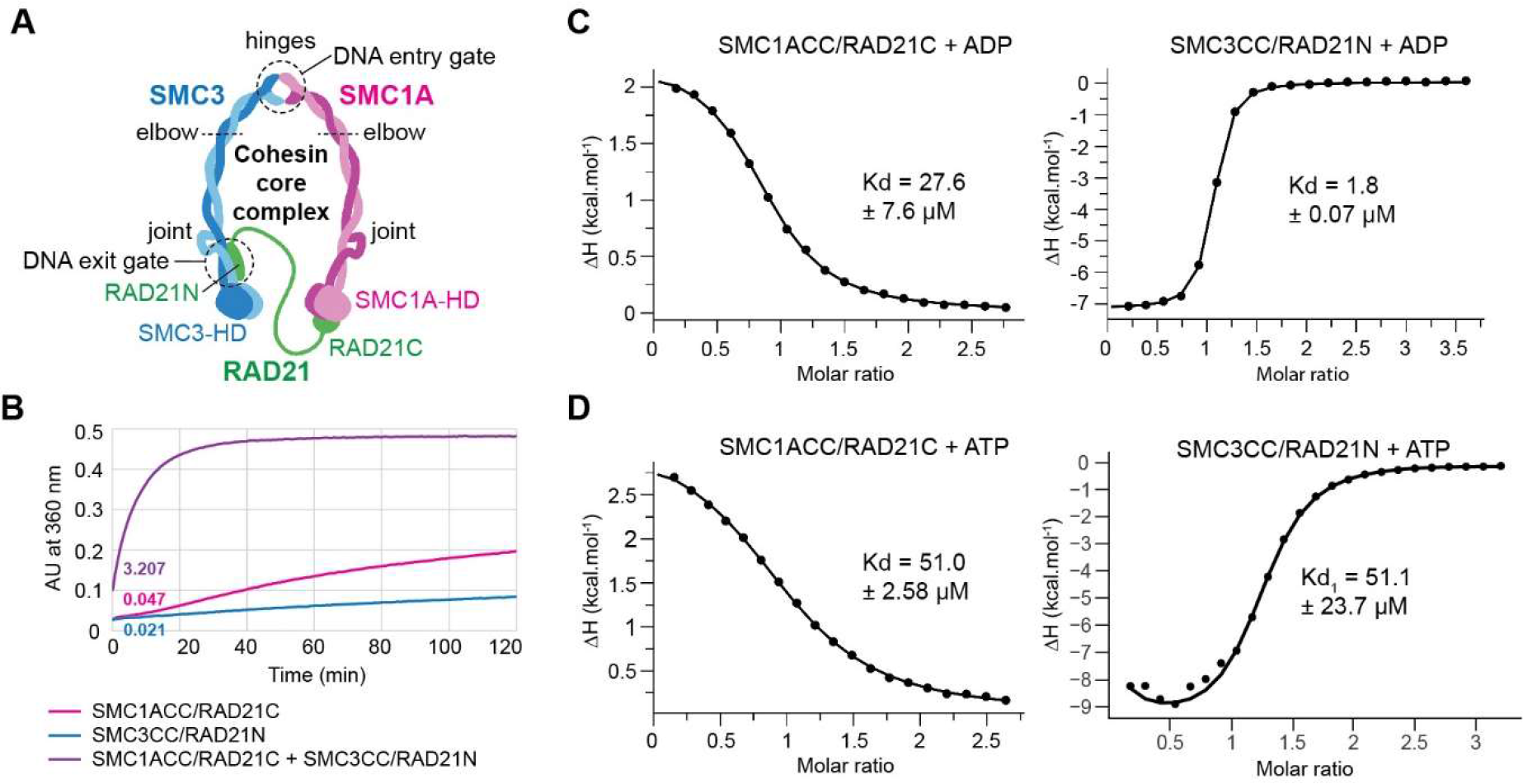
ATP hydrolysis and nucleotide binding properties of the human SMC1A and SMC3 ATPase head domains. A. Schematic representation of the human Cohesin core complex. Specific elements discussed in the text are labeled. **B.** ATPase activity of the independent and mixed SMC1ACC/RAD21C and SMC3CC/RAD21N complexes. While the independent complexes display very little ATPase activity, mixing of both complexes leads to a significant increase in ATPase activity that is similar to the ATPase activity measured in previous studies for the full-length Cohesin core complex. The ATPase activity is given in Pi molecules released per dimer and per minute. **C.** Measurements of the Kd of ADP for the SMC1ACC/RAD21C and SMC3CC/RAD21N complexes. **D.** Same as in (C) for ATP.

In the presence of ATP, the Cohesin SMC1A and SMC3 HDs also heterodimerize when binding two ATP molecules. ATP binding is driven in a composite manner by small ATP-binding motifs (Walker A/P-loop, Walker B, D-loop, Q-loop, R-loop, signature motif) contributed by both HDs at both ATP binding sites (Gligoris et al., 2014; Haering et al., 2004; Hopfner and Tainer, 2003). ATP-dependent HDs heterodimerization, known as engagement, results in the formation of the Cohesin ATPase module (engaged state) that dissociates upon subsequent ATP hydrolysis (Collier et al., 2020; Higashi et al., 2020; Muir et al., 2020; Shi et al., 2020). ATP binding and hydrolysis are key to most Cohesin mechanisms and functions, and many Cohesin auxiliary and regulatory subunits interact directly or in the vicinity of the non-engaged or ATP-engaged SMC1A and SMC3 HDs (Collier et al., 2020; Higashi et al., 2020; Petela et al., 2021; Shi et al., 2020).

Among the Cohesin regulatory subunits, NIPBL^Scc2^ has been shown to stimulate ATP hydrolysis by the Cohesin ATPase module, a stimulation required for Cohesin loop extrusion (Davidson et al., 2019; Kim et al., 2019; Murayama and Uhlmann, 2014; Petela et al., 2018). The structures of yeast and human Cohesin ATPase modules in complex with NIPBL^Scc2^ and DNA have revealed how the protein components encircle and clamp the non-topologically bound DNA (clamped state) (Collier et al., 2020; Higashi et al., 2020; Shi et al., 2020). However, by which mechanisms NIPBL^Scc2^ stimulates the Cohesin ATPase activity is still unclear. NIPBL^Scc2^ also plays an initial role in Cohesin DNA tethering by causing the ATP-dependent topological entrapment of DNA within the Cohesin ring through the hinge/hinge interface (DNA entry gate; Figure 1A) (Arumugam et al., 2003; Buheitel and Stemmann, 2013; Gruber et al., 2006; Haering et al., 2008; Higashi et al., 2020; Hu et al., 2011; Ladurner et al., 2014; Murayama and Uhlmann, 2014; Srinivasan et al., 2018).

The mechanisms subsequently leading to Cohesin genomic stabilization and then to genomic removal also require ATP binding and hydrolysis but are independent of NIPBL^Scc2^ and rely on its PDS5 regulatory subunit. Specifically, Cohesin genomic stabilization requires its association with PDS5, which is favored by the acetylation of the SMC3 HD lysines K105 and K106 and is stabilized by SORORIN (Beckouet et al., 2016; Chan et al., 2013; Chan et al., 2012; Ladurner et al., 2014; Ladurner et al., 2016; Lafont et al., 2010; Nishiyama et al., 2010; Ouyang et al., 2016; Rolef Ben-Shahar et al., 2008; Rowland et al., 2009; Unal et al., 2008; Zhang et al., 2008). In contrast, Cohesin genomic removal is also carried out by PDS5 but when bound to WAPL, DNA release occurring through opening of the SMC3/RAD21 interface (DNA exit gate; Figure 1A) (Beckouet et al., 2016; Bernard et al., 2008; Buheitel and Stemmann, 2013; Chan et al., 2012; Eichinger et al., 2013; Gandhi et al., 2006; Huis in ’t Veld et al., 2014; Kueng et al., 2006; Murayama and Uhlmann, 2015; Ouyang et al., 2016; Sutani et al., 2009; Tedeschi et al., 2013). The different mechanisms participating to Cohesin DNA tethering and release remain poorly characterized at the molecular level but are equally essential to Cohesin functions and act in synergy with loop extrusion to regulate the length of chromatin loops and to shape the vertebrate 3D genome in an ATP-dependent manner (Dauban et al., 2020; Haarhuis et al., 2017; Li et al., 2020; Nora et al., 2020; Tedeschi et al., 2013; Vian et al., 2018; Wutz et al., 2017).

An essential characteristic of Cohesin is its intrinsic flexibility, which is required for its transactions with DNA and for its biological functions. Specifically, Cohesin can adopt open ring, rod and kinked conformations depending on whether or not the SMC1A and SMC3 coiled coils are interacting by co-alignment and are bent at their elbow elements (Anderson et al., 2002; Bauer et al., 2021; Burmann et al., 2019; Chapard et al., 2019; Collier et al., 2020; Higashi et al., 2020; Huis in ’t Veld et al., 2014; Petela et al., 2021; Shi et al., 2020). The mechanisms by which Cohesin alternates between these different conformations are, however, unclear. The Cohesin ATPase cycle is expected to be key to these conformational changes. However, the specific structural rearrangements occurring within the Cohesin SMC1A and SMC3 HDs and ATPase module during the Cohesin ATPase cycle and how these can be propagated at long distance remain poorly characterized. Specifically, the structural characterization at high resolution of the complete Cohesin ATPase module, which is central to the Cohesin ATPase cycle, in the absence of any regulator has not been reported so far.

Therefore, while the Cohesin ATPase activity is at the heart of the mode of action and functions of this SMC complex, the mechanisms that underly this activity remain unclear. Here, we have characterized distinct steps of the human Cohesin ATPase cycle, including the structural determination of the Cohesin ATPase module. We show that the SMC1A and SMC3 ATPase head domains undergo highly specific structural movements and conformational rearrangements upon ATP binding, ATP-dependent engagement, ATP hydrolysis and opening of the SMC3/RAD21 interface. These movements and rearrangements are different in both HDs but concerted between them. We notably reveal the intrinsic flexibility of the SMC3 HD proximal coiled coil and the high plasticity of the ATP-dependent interface between the SMC1A and SMC3 HDs that underlie the conformational dynamics of these HDs. Our results show the importance of the full N-terminal region of RAD21 in these mechanisms, which we confirm *in vivo* using the zebrafish. Collectively, our results provide a most complete view of the Cohesin ATPase cycle and of its specific conformational dynamics that support Cohesin mechanisms and functions.

## Results

### The human SMC1A and SMC3 ATPase HDs have distinct nucleotide binding properties and can recapitulate the ATPase activity of the full core Cohesin complex

The characterization of the Cohesin complex in molecular terms is rendered difficult by the large flexibility of this complex. We have used the isolated SMC1A and SMC3 ATPase HDs to obtain the high-resolution data required to study in precise molecular terms the conformational dynamics of these HDs. Different constructs of the human SMC1A and SMC3 HDs were made by varying the length of the proximal coiled coil (CC) emerging from their globular domains (GD): short (SMC1ACCsh), up to the joint element (SMC1ACC and SMC3CC), and including the joint element (SMC1AJ and SMC3J) (Figure 1A; Supplementary Figure 1A,B). The RAD21N and RAD21C constructs were designed according to previous work (Supplementary Figure 1C) (Gligoris et al., 2014; Haering et al., 2004). While single expression of these constructs led to insoluble proteins, their co-expression led to the soluble production of the various SMC1A HD/RAD21C and SMC3 HD/RAD21N complexes, with the exception of the SMC1AJ/RAD21C complex that was poorly soluble. The SMC1ACC/RAD21C and SMC3CC/RAD21N complexes were chosen as primary targets for our investigations unless stated.

ATPase activity measurements showed that the SMC1A HD/RAD21C and SMC3 HD/RAD21N complexes have a very low ATPase activity when analysed independently (Figure 1B). However, upon mixing of these complexes, a significant increase in ATPase activity was observed (Figure 1B). Importantly, the ATPase activity of 3.2 Pi molecules released per dimer and per minute that we measured for the mixed ATPase HDs is within the range (∼2.4-4.0) measured for the full-length human core Cohesin complex obtained either by co-expression and purification using the baculovirus expression system or endogenously purified from HeLa cells (Davidson et al., 2019; Kim et al., 2019; Ladurner et al., 2014). This demonstrates that the isolated HDs used for our study can recapitulate the ATPase activity of the full core Cohesin complex.

Measurement using ITC of the Kd of nucleotides for the individual SMC1ACC/RAD21C and SMC3CC/RAD21N complexes was performed both with the wild-type (WT) and with the low ATPase activity EQ mutants (SMC1A-E1157Q and SMC3-E1144Q) as well as with ATP and the low hydrolysable ATPγS analogue. While the measured Kd values were all in the µM range, varying however depending on the nucleotide analysed (Figure 1C-D; Supplementary Figure 2A-B; Supplementary Table 1), significant differences were observed for the ITC-measured thermodynamic signatures of both complexes. Specifically, nucleotide binding to SMC1ACC/RAD21C was entropy-driven, suggesting a solvent entropy increase, whereas binding to SMC3CC/RAD21N was enthalpy-driven (Supplementary Figure 2C-D).

We next used X-ray crystallography to investigate the molecular properties of the independent SMC1ACC/RAD21C and SMC3CC/RAD21N complexes and their binding to nucleotides. Crystallizations assays were performed without nucleotide or in the presence of either ADP or ATPγS. All complexes crystallized in apo form and in complex with ADP or ATPγS. The crystals obtained diffracted between 1.4 and 3.1 Å resolution (Supplementary Tables 2, 3 and 4). Only the crystals of the SMC3J/RAD21N complex showed low diffraction and were not considered further. Structure determinations were carried out by molecular replacement.

### The SMC1A HD/RAD21C complex shows Q-loop-based nucleotide-induced rotational movements but remains in a specific relaxed conformation

Analysis of the SMC1A HD/RAD21C structures showed that the SMC1A HD P-loop adopts, in the absence of nucleotide, a closed conformation that is stabilized by a network of water molecules interacting with neighbouring residues (Supplementary Figure 3A-B). The movement of the P-loop from a closed to an open conformation upon nucleotide binding therefore requires the release of this water network, which agrees with the observed entropy-driven binding of ADP and ATP to SMC1ACC/RAD21C. In addition, in all our structures, R-loop R57 is stably interacting with N40 and D43 and, in the presence of ATPγS, with the α-phosphate group of this nucleotide (Supplementary Figure 3C-D). This contrasts with fungi where the Smc1 R-loop is not defined in density unless in the clamped state (Collier et al., 2020; Haering et al., 2004; Muir et al., 2020). Mutation of R57 into alanine (R57A) or the use of the Cornelia de Lange Syndrome (CdLS)-characterized 58-62 deletion mutant (Δ58-62) (Rohatgi et al., 2010) caused a complete loss of ATP hydrolysis by the SMC1A HD (Supplementary Figure 3E). However, this loss was only ∼10% when SMC3CC/RAD21N was added to the reaction (Supplementary Figure 3E). This indicates that human SMC1A R57 plays an early role in organizing the SMC1A HD nucleotide binding site prior to HDs engagement.

Comparative analysis of the various SMC1ACC/RAD21C structures revealed specific conformational changes occurring upon ADP and ATPγS binding. In both cases, reorganizations in the RecA-lobe to adapt to the bound nucleotide induce a slight rotational movement toward the Helical-lobe of the region of the RecA-lobe that interacts with the adenine moiety of the nucleotide. In the case of ADP binding, this causes the Helical-lobe to rotate away from the RecA-lobe (Figure 2A). In contrast, ATPγS binding causes the Helical-lobe to rotate toward the RecA-lobe (Figure 2B). These rotational movements occur in a same plan that goes through the Q-loop (Figure 2C-D). Notably, upon ATPγS binding, Q137 moves toward the nucleotide binding site and binds to the Mg ion (1.9 Å) and hydrogen bonds to the γ-thiophosphate group (2.6 Å) (Figure 2E). These movements do not induce significant structural changes within the Helical-lobe itself and to the RecA-lobe/Helical-lobe interface but position differently the SMC1A proximal coiled coil depending on the nucleotide bound (Figure 2C).

**Figure 2.**
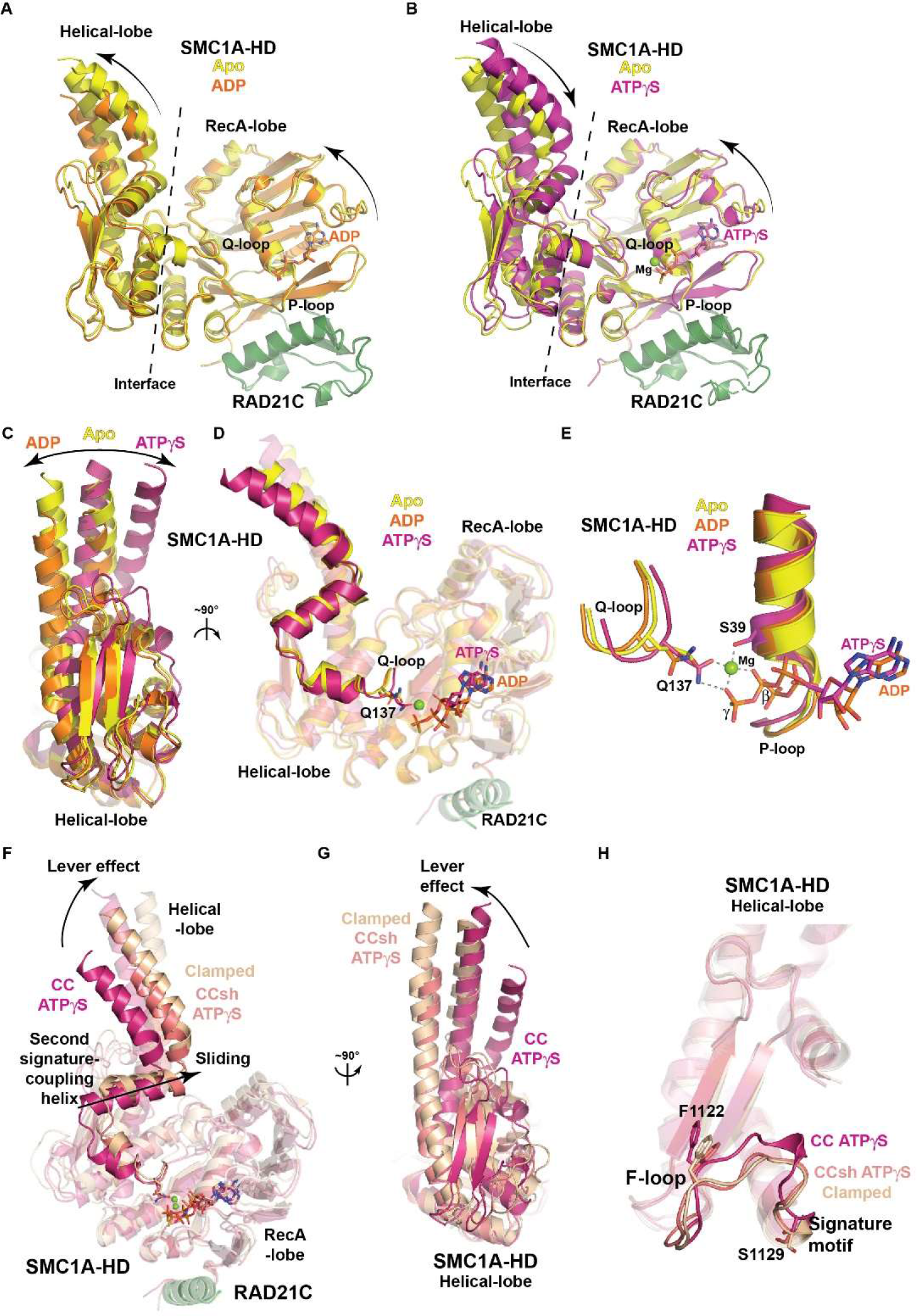
Specific conformational dynamics of the SMC1A HD. A. Conformational movements of the SMC1A HD upon ADP binding. ADP binding causes rotational movements in the same direction of the RecA-lobe and Helical-lobe. **B.** Same as in (A) upon ATPγS and magnesium binding. In contrast to ADP binding, the RecA-lobe and Helical lobe of the SMC1A HD rotate in opposite directions upon ATPγS binding, bringing the SMC1A CC closer to the RecA-lobe. **C.** Opposite rotational movements of the SMC1A Helical-lobe upon ADP and ATPγS binding. **D.** The movements observed in (D) are in a same plane that passes through Q137. **E.** Close-up of (D) on the Q-loop. Specific movements of the SMC1A Q-loop and Q137 depend on the bound nucleotide. Upon ATPγS and magnesium binding, Q137 moves toward the ATP binding site, hydrogen bonds with the γ- thiophosphate group of the ATPγS molecule and interacts with the magnesium ion. **F.** Movements of the Helical-lobe with respect to the RecA-lobe of SMC1A upon formation of the clamped complex. These movements are caused by a lever effect which induces a sliding along the RecA-lobe of the second signature-coupling helix of SMC1A and a major displacement of the SMC1A CC. These movements are different from the rotational movements observed upon nucleotide binding by the independent SMC1A HD. Therefore, when non-engaged, the SMC1A HD adopts a relaxed conformation, regardless of its nucleotide-binding state. A similar, artificially-induced lever effect is observed in the SMC1ACCsh/RAD21C structures. **G.** Same as in (A) in a 90° view focusing on the SMC1A Helical-lobe. A repositioning of the SMC1A CC like in the apo form is observed upon the lever effect caused by the formation of the clamped complex. **H.** Conformational and positional changes occurring in the SMC1A F-loop and signature motif upon formation of the clamped complex. These are the only significant changes occurring in the helical lobe and are linked to the reorganization of the RecA-Lobe/Helical-lobe interface.

Comparison of our SMC1ACC/RAD21C structures with that of the SMC1A HD in the clamped state (Shi et al., 2020) revealed major conformational changes occurring within the SMC1A HD upon clamping. Notably, a large movement of the SMC1A HD Helical-lobe with respect to the RecA-lobe is observed upon formation of the clamped state (Figure 2F). This movement, which has been termed lever effect in the bacterial/archaeal case (Kamada et al., 2017), is due to a sliding of the Helical-lobe along the RecA-lobe. The amplitude of this movement is large, as demonstrated by the displacement by ∼7-8 Å of the second signature-coupling helix of SMC1A that precedes the N-terminal coil of this HD (Figure 2F). The lever effect also repositions the SMC1A CC in a central position (Figure 2G), as observed in our apo structure, but keeps Q137 at a close distance to the ATP molecule (Shi et al., 2020).

While the overall structure of the SMC1A Helical-lobe is not significantly modified by the lever effect, the F-loop and signature motif of this lobe nevertheless undergo significant conformational and positional changes upon formation of the clamped complex (Figure 2H). This also requires a significant reorganization of the RecA-lobe/Helical-lobe interface. Importantly, the sliding movement upon leverage is almost perpendicular to the aforementioned planar rotational movement observed upon ADP and ATPγS binding to the SMC1A HD, showing that these conformational changes are not related (Figures 2D,F). This shows that, on its own, the SMC1A HD adopts a stable relaxed conformation, regardless of its nucleotide-binding state, which is different from that of the clamped state.

Of note, our crystallographic structures of the SMC1ACCsh/RAD21C complex, where the SMC1A HD proximal coiled coil is shorter, show a higher structural resemblance with that of the clamped SMC1A HD. These structures notably display, as in the clamped conformation, a lever effect, a central repositioning of the SMC1A CC, a specific RecA-lobe/Helical-lobe interface and a conformational rearrangement of the F-loop and signature motif compared to the SMC1ACC/RAD21C structures (Figure 2F-H). These features appear, however, artificially induced by the crystal packing of the SMC1ACCsh/RAD21C complex that involves the SMC1A CC but not the ATP binding interface (Supplementary Figure 4). This shows that the reorganization of the SMC1A HD F-loop only requires the rearrangement of the RecA-lobe/Helical-lobe interface.

### The SMC3 HD/RAD21N complex adopts a specific resting state through interactions of RAD21N with the SMC3 HD globular domain

In contrast, our SMC3CC/RAD21N structures show a higher structural similarity whatever their nucleotide binding state, with only limited adaptative movements observed upon binding of ADP and ATPγS (Figure 3A). In addition, as previously observed in the clamped state, the R-loop, including R57, and the two K105 and K106 lysines, which are targets of the ESCO1/2^Eco1^ acetylases, are exposed at the surface of the SMC3 GD and poised for interaction with incoming macromolecules (Collier et al., 2020; Gligoris et al., 2014; Higashi et al., 2020; Shi et al., 2020) (Figure 3A). More surprising, in all our structures the SMC3 Q-loop is positioned far away (∼9 Å) from the nucleotide binding site, with the Q141 side chain turned away from this binding site, even when ATPγS and magnesium are bound (Figure 3B).

**Figure 3.**
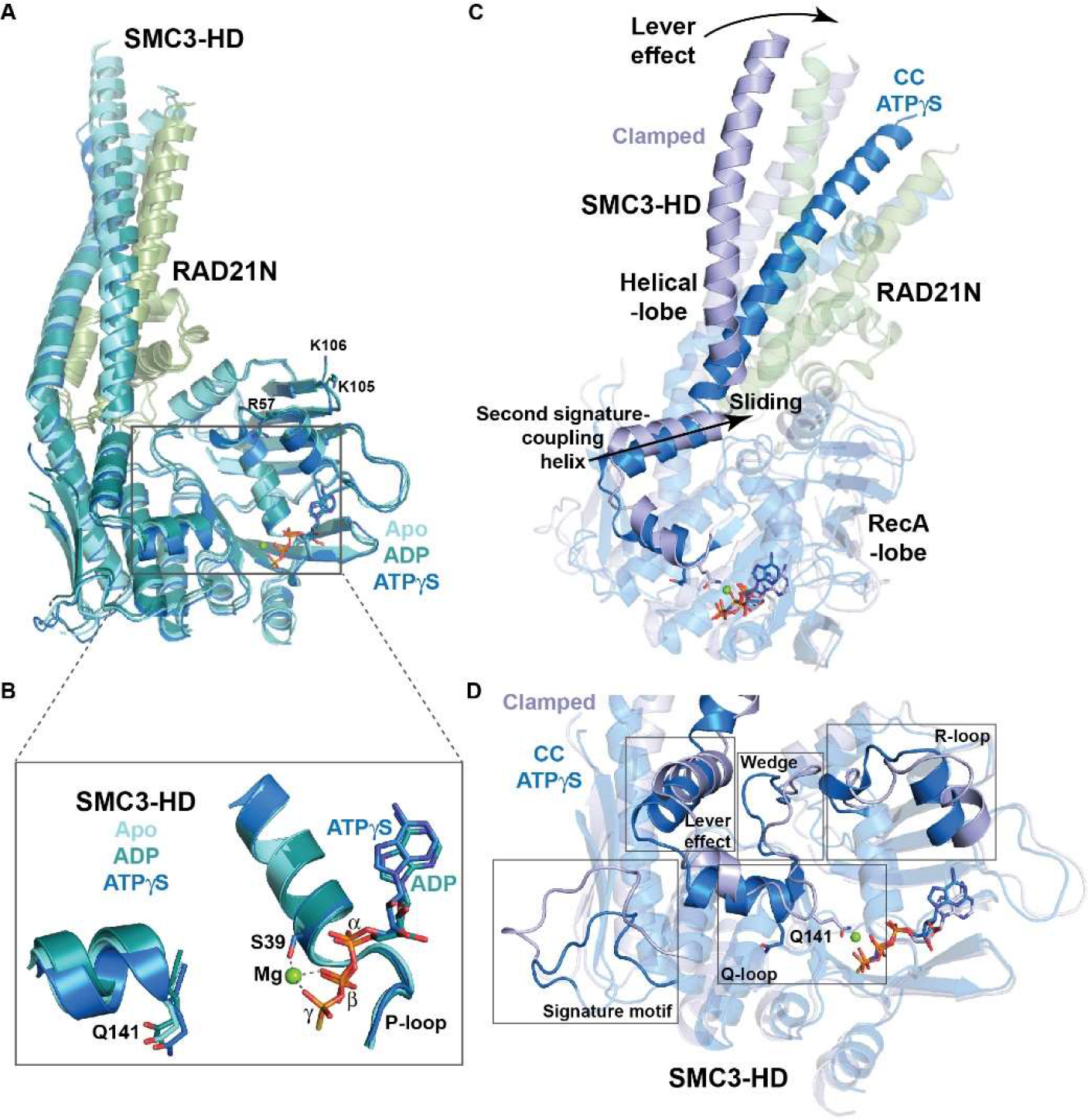
Specific conformational dynamics of the SMC3 HD. A. Superposition of the SMC3CC/RAD21N structures in various nucleotide-binding states (apo, ADP-bound and ATPγS-bound) showing that, in contrast to the SMC1A HD, the SMC3 HD adopts a stable conformation regardless of its nucleotide-binding state. **B.** Close-up on the ATP binding site of the structures shown in (A). Q141 remains turned away from the active site even in the presence of ATPγS and magnesium. **C.** Conformational changes of the SMC3 HD upon formation of the clamped complex, including, like for the SMC1A HD, a lever effect that slides the second signature-coupling helix of this HD along the RecA-lobe. **D.** Multiple specific changes in the SMC3 HD induced by the formation of the clamped complex.

Comparative analysis shows that, to reach the clamped conformation, the SMC3 HD also requires a lever effect that slides its Helical-lobe along its RecA-lobe, moving the SMC3 second signature-coupling helix by ∼4-5 Å (Figure 3C-D). In this case, the associated conformational changes affect the positioning and conformation of the Q-loop, and reposition Q141 toward the ATP-binding site (Figure 3C-D). Additional changes, specific to the SMC3 HD, are also observed that are reminiscent of those observed upon bacterial/archaeal Smc HD homodimerization (Kamada et al., 2017). Notably, the formation of the clamped complex modifies the conformation of the wedge element, which turns away from the SMC3 CC toward the R-loop, accompanying the repositioning and conformational change of this loop (Figure 3D). In addition, a repositioning but not a conformational rearrangement of the loop harbouring the SMC3 signature motif is observed (Figure 3D).

The most striking difference observed, however, is the ∼45° kink of the CC emerging from the SMC3 globular domain (GD) in our structures compared to the straight conformation displayed by this CC in the clamped complex (Figure 4A-C). This kink is due to a simultaneous bending of both α-helices forming the human SMC3 CC that occurs right above the region where these helices emerge from the SMC3 GD, and just below the region where they interact with RAD21N (Figure 4A-B). The intrinsic flexibility of the short helical stretches of the SMC3 CC that enables their bending is demonstrated by their slightly different conformations observed in our different structures, including between non-crystallographic symmetry mates in a single crystal form.

**Figure 4.**
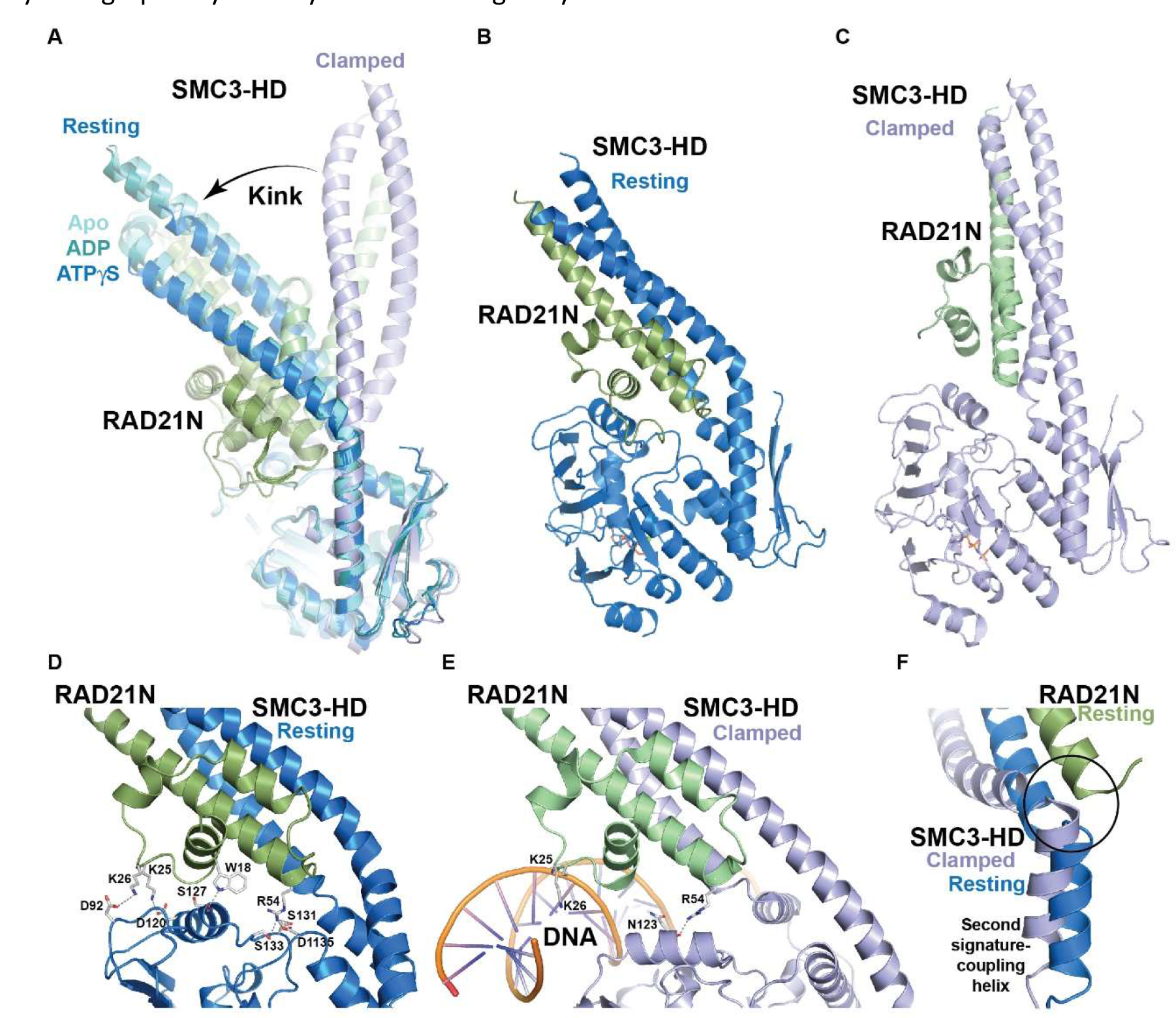
The independent SMC3 HD/RAD21N complex adopts a stable resting state. A. Comparison of the structures of the independent SMC3 HD/RAD21N complex with that of this complex in the clamped conformation. A major kink of the SMC3 CC is observed in the independent structures that repositions RAD21N without perturbing the structure and integrity of the SMC3 CC/RAD21N interaction. This resting state of the SMC3 HD/RAD21N complex is enabled by the inherent flexibility of short stretches of both coils in the SMC3 CC. **B.** Structure of the SMC3 HD/RAD21N complex in its resting conformation showing the positioning of the RAD21N HelD domain at the interface between the RecA-lobe and Helical-lobe of the SMC3 HD. **C.** Structure of the SMC3 HD/RAD21N complex in its clamped conformation showing that RAD21N is positioned further away and hardly interacts with the SMC3 GD. **D.** Interactions made by the RAD21N HelD domain with the SMC3 GD in the resting state. **E.** Interactions made by the RAD21N HelD domain with the SMC3 GD and the DNA in the clamped complex. **F.** Steric clashes (circled) between the SMC3 second signature-coupling helix in the clamped conformation and the RAD21N long α-helix paralleling the SMC3 CC in the resting state. The resting conformation of the SMC3 HD/RAD21N complex is incompatible with the lever effect in the SMC3 HD.

The SMC3 CC kink is observed in all our SMC3CC/RAD21N structures, regardless of the nucleotide-binding state. Although the folding of RAD21N and its interaction with the SMC3 CC are not changed by the kink, the new path of the SMC3 CC positions differently the RAD21 50-residues small N-terminal helical domain (HelD), which precedes the long RAD21N α-helix interacting with the SMC3 CC, in the groove at the interface between the RecA-lobe and Helical-lobe of the SMC3 GD (Figure 4B). The interactions formed between the RAD21N HelD and the SMC3 GD stabilize this specific conformation of the SMC3 HD/RAD21N interface that we have termed resting state (Figure 4D).

Specifically, in the resting state, RAD21N residues 5 to 26 pack against the SMC3 GD forming hydrophobic and hydrogen bonding interactions. Notably, the first small α-helix of the RAD21N HelD, composed of residues 13 to 23, packs against the SMC3 GD, including tryptophan W18 whose indole nitrogen makes a hydrogen bond with the main chain carbonyl of SMC3 S127. Following this helix, RAD21N K25 and K26 side chains interact respectively with the SMC3 GD D120 and D92 side chains. In addition, the R54 guanidino group makes several interactions with the SMC3 GD, notably with the acidic patch formed by the side chains of S131, S133 and D1135 (Figure 4D).

In comparison, in the clamped state, the RAD21N HelD is positioned above the SMC3 GD, both domains only forming a single contact through the interaction established between the RAD21N R54 side chain and the SMC3 N123 main chain carbonyl (Figure 4E) (Shi et al., 2020). In addition, the clamped RAD21N HelD interacts with the bound DNA through electrostatic interactions and through direct contacts, such as those that can be formed by RAD21N K25 and K26 with the DNA phosphate backbone (Figure 4E). Importantly, the position of RAD21N in the resting state, notably its long α-helix interacting with the SMC3 CC, is not compatible with the forward movement of the SMC3 second signature-coupling helix that occurs upon leverage (Figure 4F). Therefore, SMC3 is kept in its resting conformation through stabilization of the RAD21N HelD onto the SMC3 GD, thereby hampering the structural rearrangements required to reach the clamped state.

### ATP-dependent engagement of the SMC1A and SMC3 HDs induces large conformational changes that do not fully recapitulate those observed in the clamped state

To further characterize the Cohesin ATPase cycle, we next investigated the conformational dynamics of the SMC1A and SMC3 HDs upon their ATP-dependent engagement to understand the molecular mechanisms leading to the release of the respective relaxed and resting conformations of the SMC1A and SMC3 ATPase HDs. We first looked at conditions causing stable HDs interaction by using size exclusion chromatography since weakly formed complexes cannot resist the dilution conditions of this technique. Analytical ultracentrifugation was used to confirm the results obtained.

Initial analyses with the WT and EQ constructs revealed that only the SMC1ACC-EQ/RAD21C and the SMC3CC-EQ/RAD21N mutant complexes can stably associate when ATP is present (Figure 5A-B).

**Figure 5.**
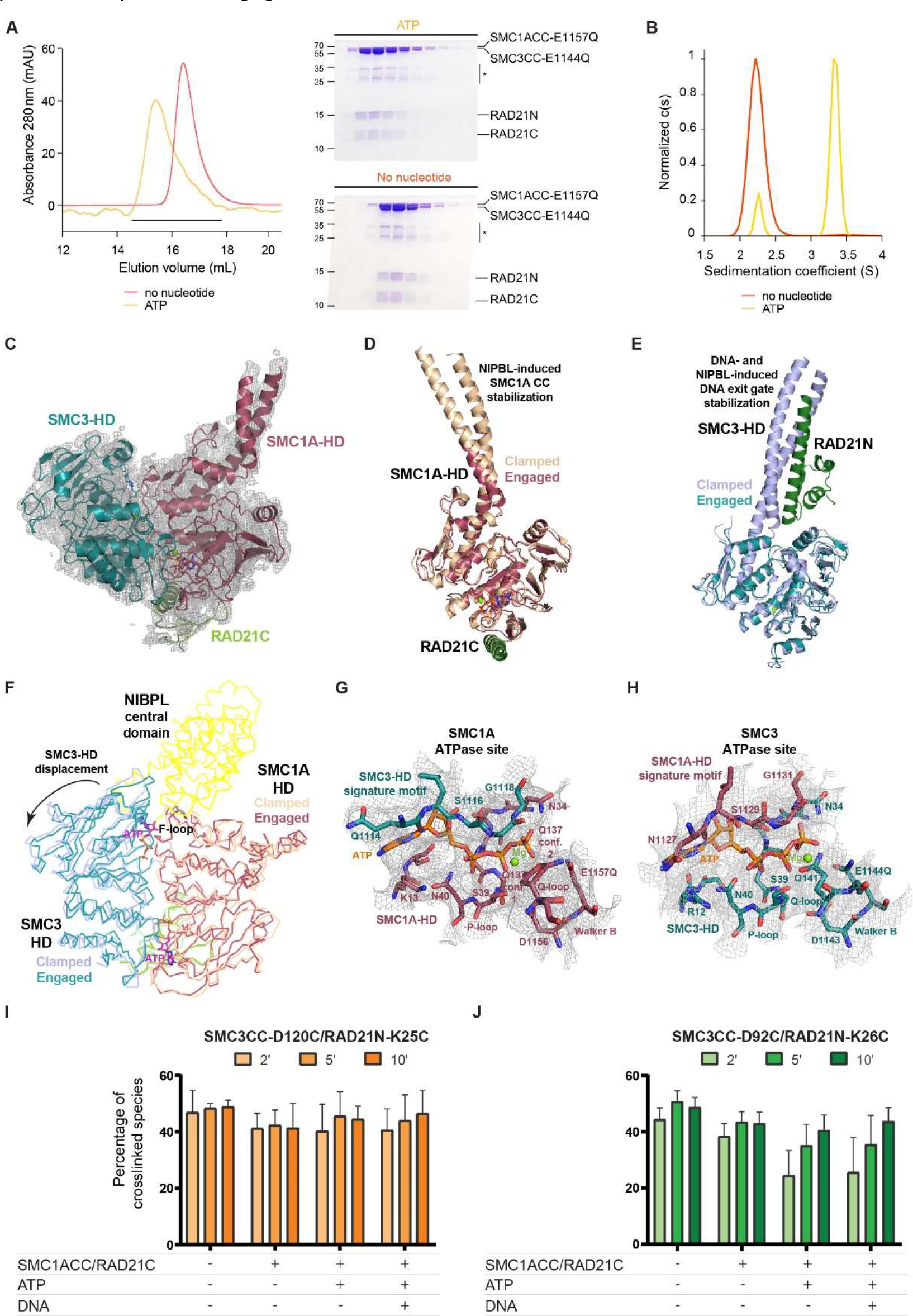
ATP-dependent engagement of the SMC1A and SMC3 HDs yields an engaged ATPase module with a flexible SMC3 CC/RAD21N complex. A. Size exclusion chromatographic profiles and associated SDS-PAGE analyses of the mixed SMC1ACC- EQ/RAD21C and SMC3CC-EQ/RAD21N mutants in the absence (red) and the presence (orange) of ATP. In the presence of ATP, a shift of the peak toward the high molecular weight fractions demonstrates of the stable ATP-dependent engagement of the HDs to form the Cohesin ATPase module. The fractions analysed by SDS-PAGE are indicated with a black bar in the chromatographic profiles. **B.** Analytical ultracentrifugation velocity experiments confirming that the HDs used in (A) can form a stable Cohesin ATPase module in the presence of ATP. **C.** Structural model of the human engaged ATPase module displayed within the 3.6 Å resolution cryo-EM map. The map shows the absence of a conformationally stable SMC3 CC/RAD21N complex, implying that, upon engagement, this complex is released from its resting conformation. **D.** Superposition of the SMC1A HDs of the engaged and clamped ATPase modules. Both structures display a high similarity, showing that engagement already induces most of the conformational changes observed for this HD in the clamped complex. Nevertheless, in the clamped state, a stabilization occurs of the upper region of the SMC1A CC that interacts with the C-terminal part of the hook domain of NIPBL^Scc2^. **E.** Superposition of the SMC3 HDs of the engaged and clamped ATPase modules. Both structures display a high similarity, showing that engagement also induces most conformational changes observed in the clamped complex. The SMC3 CC/RAD21N complex is, however, not seen in the cryo-EM map of the engaged ATPase module. For clarity, the SMC3 Joint element in the clamped complex is not shown. **F.** Superposition of the engaged and clamped ATPase modules using the SMC1A HD as reference. A displacement of the SMC3 HD with respect to the SMC1A HD is observed between both structures that appears due to the binding of the central region of NIPBL^Scc2^ (yellow) to the SMC1A F-loop in the groove at the edge of the SMC1A/SMC3 interface. For clarity, only the region of NIPBL^Scc2^ binding to the F-loop is displayed. **G.** Representation of the ATP-binding motifs forming the SMC1A composite ATPase active site and of the bound ATP molecule and magnesium ion displayed in the 3.6 Å cryo-EM map. The SMC1A Q137 side chain appears to adopt two conformations, one turned toward the ATPase site and the other turned away toward E1157Q. **H.** Same as in (G) for the SMC3 composite ATPase site. The positioning of the ATP γ-phosphate group appears less constrained. **I.** Quantification of crosslinked species for the SMC3CC-D120C/RAD21N-K25C pair in experiments performed at room temperature. The supplementation of the SMC1ACC/RAD21C, ATP and DNA is indicated underneath the graph. All experiments were done in triplicates. **J.** Same as in (I) for the SMC3CC-D192C/RAD21N-K26C pair. The longer distance between the SMC3 GD and RAD21N in the clamped state enables the observation in a time-dependent manner of the increase of crosslinking that indicates the oscillation of the SMC3 CC/RAD21N complex upon engagement. DNA binding does not alter significantly this dynamic.

Injection of the two EQ mutants on the ATP-loaded chromatographic column without preincubation with ATP was sufficient to form the complex, demonstrating its rapid assembly and stability. In contrast, combination of WT/WT, WT/EQ and EQ/WT complexes failed to provide a stably engaged ATPase module in the presence of ATP (Supplementary Figure 5A). The fact that RAD21N was not lost during these experiments in diluting conditions shows that the SMC3 HD/RAD21N interface remains shut upon ATP-dependent engagement.

In addition, both size exclusion chromatography and analytical ultracentrifugation showed that while ATP causes engagement, ADP, ATPγS and AMP-PNP cannot (Supplementary Figure 5B), in agreement with previous observations on the yeast ATPase module (Hu et al., 2011). Moreover, we did not observe a homodimerization of the independent EQ mutants in the presence of ATP, demonstrating of the specificity of the heterodimeric engagement (Supplementary Figure 5C). Mutants in the signature motifs (SMC1A L1128V and SMC3 L1115V) and D-loops (SMC1A D1163E and SMC3 D1150E) have previously been shown, like the EQ mutants, to have a low ATPase activity and to enable engagement (Beckouet et al., 2016; Camdere et al., 2015; Elbatsh et al., 2016). However, no combination of these mutants yielded stable engagement in our experimental conditions (Supplementary Figure 5D).

We used this stable engaged ATPase module for structural analyses. Crystallization attempts of the complex only yielded poorly diffracting crystals and cryo-electron microscopy was used instead using the purified samples obtained directly from the size exclusion chromatography without further treatment. Processing of the collected data yielded two cryo-EM maps at 3.6 and 4.4 Å resolution, both representing engaged complexes (Supplementary Figure 6). The best resolved 3.6 Å map was used to solve the structure of the engaged human Cohesin ATPase module. We established a first model of this module by fitting our high-resolution crystallographic structures into this map. This model was further refined by cycles of manual building and automatic real space refinement (Figure 5C and Supplementary Table 5).

The refined structure of the engaged human Cohesin ATPase module shows strong similarities to that in the clamped state. Notably, upon engagement, both SMC1A and SMC3 HDs undergo lever effects that release them from their specific relaxed and resting conformations (Figure 5D-E). In addition, engagement repositions the SMC1A F-loop, the SMC1A and SMC3 signature motifs, and the SMC3 wedge element and R-loop similarly as in the clamped state. Therefore, several of the structural rearrangements observed in the ATPase module in the clamped state are already occurring at the engagement stage.

However, despite these similarities, the structures of the engaged and clamped ATPase modules also show major differences. Notably, comparison of the heterodimerized ATPase modules reveals that, while ATP binding is retained at both ATPase sites, the interaction between both HDs is tighter in the engaged complex than in the clamped complex. Specifically, the superposition of these complexes using the SMC1A HD for fitting, reveals a displacement by 2-3 angstroms of the SMC3 HD when comparing the engaged complex with the clamped complex (Figure 5F). This appears due to the binding of the central domain of NIPBL^Scc2^ to the SMC1A F-loop in the clamped state that causes steric hindrances at the interface between both HDs that cannot be accommodated solely by local rearrangements (Figure 5F).

The observed displacement does not cause loss of ATP at both ATPase sites and is enabled by significant changes within the complete SMC1A HD/SMC3 HD interface, including the repositioning with respect to each other of the composite ATP binding/hydrolysis motifs forming both sites. Specifically, in the engaged complex, the positioning of these motifs appears to enable SMC1A Q-loop Q137 side chain to adopt two different rotamer conformations, either turned toward the ATP molecule and the magnesium ion or toward the SMC1A E1157Q residue (Figure 5G). In the case of SMC3, while the Q141 side chain is now turned toward the ATPase site, the ATP γ-phosphate group appears imperfectly positioned to interact stably with this residue due to a void created between the ATP-binding motifs and within which the γ-phosphate of the bound ATP molecule can move (Figure 5H). These features could potentially contribute to the observed low ATPase activity of the engaged ATPase module.

### Engagement induces an oscillation of the SMC3 CC/RAD21N complex

Our structural comparison between the engaged and clamped complexes also reveals major differences for the SMC1A and SMC3 CCs. In the case of the SMC1A HD, 3-5 helical turns within each helix of its CC are stabilized in the clamped complex, whereas these are otherwise disordered in our cryo-EM map of the engaged ATPase module (Figure 5D). This region of the SMC1A CC is that interacting directly with NIPBL^Scc2^ in the clamped complex. The difference is even larger in the case of the SMC3 HD since, in our cryo-EM map, there is a complete absence of density for the SMC3 CC/RAD21N complex. Specifically, the density for the SMC3 CC is lost where this CC is kinked in our crystallographic structures (Figure 5C,E).

The absence of a resting conformation for the SMC3 CC/RAD21N complex in the engaged ATPase module agrees with the fact that the lever effect is incompatible with the resting state of the SMC3 HD (Figure 4F). This suggests that engagement releases the resting conformation of the SMC3 HD through the repositioning of the SMC3 second signature-coupling helix upon leverage. This release of the resting conformation does not seem, however, to cause a straightening of the SMC3 CC that would enable the SMC3 HD/RAD21N complex to adopt a conformation similar to that observed in the clamped state (Figure 5E), suggesting that this complex oscillates between these two conformations. To investigate this dynamic of the SMC3 CC/RAD21N complex upon engagement, we have performed site-specific crosslinking experiments in solution between the SMC3 GD and RAD21N. Based on our structures and the structure of the clamped complex, we identified two pairs of residues in the SMC3 GD and in RAD21N (SMC3CC-D120/RAD21N-K25 and SMC3-D92/RAD21N-K26) that could be used for these experiments (Figure 4D-E). These residues, when mutated to cysteines, are well positioned for crosslinking with Bis-maleimidoethane (BMOE) in a conformation close to the resting state (8.5 and 9.5 Å, respectively) but not in the clamped state (14.0 Å and 25.0 Å, respectively).

Crosslinking experiments with the SMC3CC-D120C/RAD21N-K25C complex gave a robust crosslinking (> 40-50%). As expected, the amounts of crosslink were not significantly modified in the presence of SMC1A alone. Additional supplementation of ATP or ATP and DNA, however, did not further affect crosslinking (Figure 5I and Supplementary Figure 7). We reasoned that this could possibly be due to the insufficient amplitude of the oscillation of the SMC3CC/RAD21N complex (maximal distance of 14 Å) that keeps the cysteine residues at crosslinking distance. Accordingly, using the SMC3- D92C/RAD21N-K26C complex, where the distance between the crosslinking residues increases more rapidly upon oscillation (maximal distance of 25.0 Å), a significant difference in crosslinking was observed in a time-dependent manner in the presence of SMC1A when supplemented with ATP or ATP and DNA (Figures 5J and Supplementary Figure 7). These results therefore confirmed that, upon ATP-dependent engagement, the resting state of the SMC3 HD/RAD21N complex is released and that the SMC3CC/RAD21N complex oscillates rather than adopting a stable conformation.

### Opening of the SMC3 HD/RAD21N interface keeps engagement but stiffens the SMC3 CC and positions the SMC1A and SMC3 CCs in the ATPase module DNA binding chamber

We then used our 4.4 Å resolution cryo-EM map for model building. Surprisingly, this map showed unambiguous density for the SMC3 CC emerging from the SMC3 GD but not for RAD21N. We therefore fitted our 3.6 Å resolution engaged ATPase module structure into this map, added the SMC3 CC and performed cycles of main chain manual modification and automatic real space refinement to obtain a structural model. This confirmed the presence of a structured SMC3 CC but not of RAD21N (Figure 6A). Due to the lack of RAD21N, we termed open-engaged this second complex of the human ATPase module. Structural comparison showed that our human open-engaged complex is similar to the yeast ATPase module lacking Scc1N where the Smc3 CC also adopts a stable conformation, thus confirming our observations (Muir et al., 2020).

**Figure 6.**
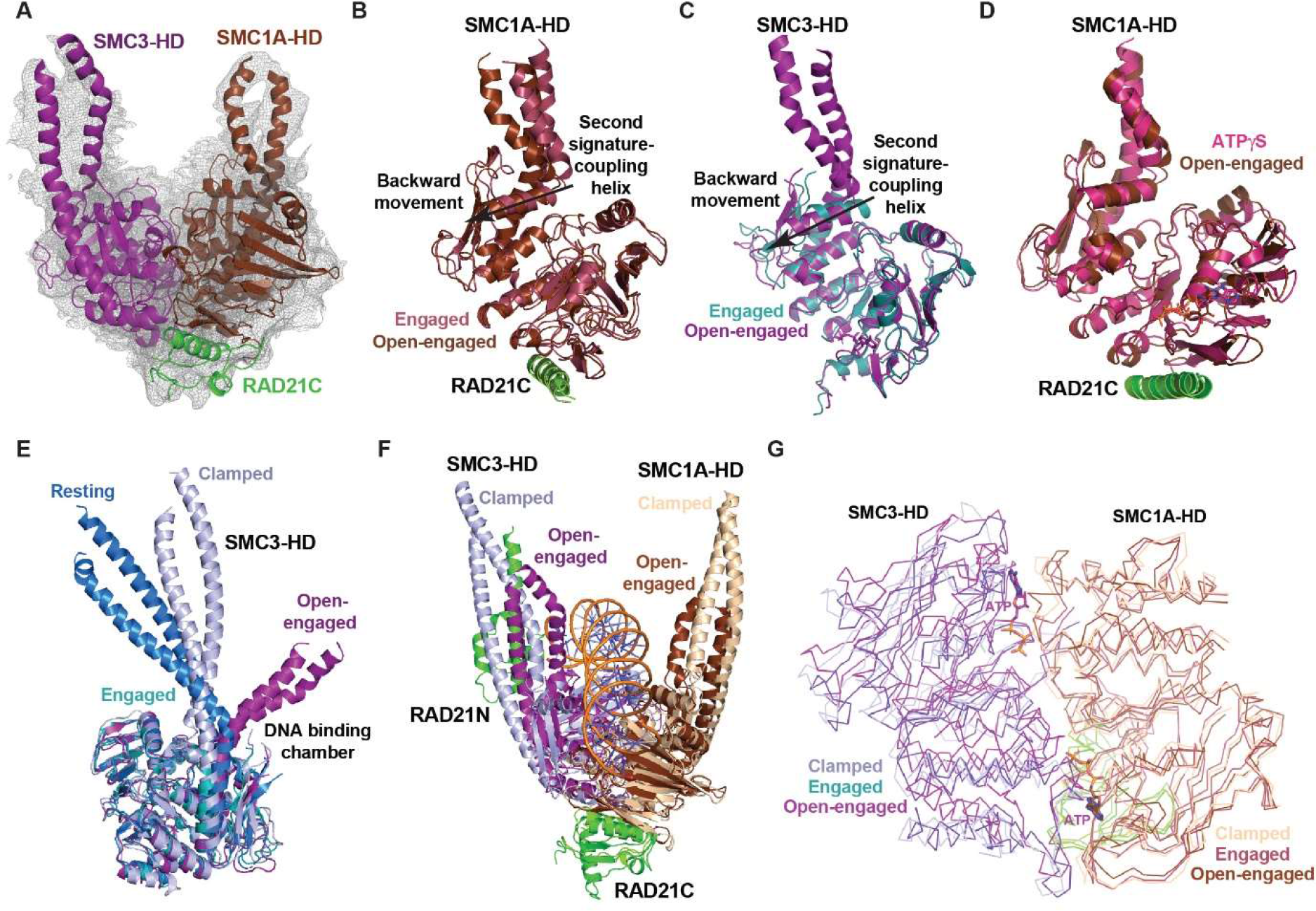
Opening of the SMC3 CC/RAD21N interface keeps the ATPase gate shut but causes the stiffening of the SMC23 CC that constricts the DNA binding chamber of the Cohesin ATPase module. A. Structural model of the human open- engaged ATPase module displayed within the 4.4 Å resolution cryo-EM map. The map shows a stabilization of the SMC3 CC emerging from the SMC3 GD. **B.** Superposition of the SMC1A HDs of the open-engaged and engaged ATPase modules. A backward lever effect movement of the Helical-lobe with respect to the RecA-lobe is observed. **C.** Same as in (B) for the SMC3 HDs. A similar backward lever effect movement is observed. **D.** Superposition of the open-engaged SMC1A HD/RAD21C complex with that of the ATPγS-bound independent SMC1A HD/RAD21C complex. Both structures show a strong similarity, including with their CCs that are positioned similarly with respect to the RecA-lobe. **E.** Structural superposition of the resting (blue), engaged (green-cyan), open-engaged (violet) and clamped (light blue) SMC3 HD structures. The SMC3 CC shows a high but specific conformational flexibility when emerging from the SMC3 GD. Notably, in its stabilized open-engaged conformation, the SMC3 CC is turned toward the ATPase module DNA binding chamber. **F.** Superposition of the clamped and open-engaged complexes. The conjoint movements of the SMC1A and SMC3 CCs upon loss of RAD21N constrict the ATPase module DNA binding chamber. **G.** Superposition of the engaged, open-engaged and clamped ATPase modules using the SMC1A HD as reference. An even larger repositioning of the open-engaged SMC3 HD with respect to the SMC1A HD is observed compared to the other complexes. The two ATP molecules driving the interaction of both HDs remain bound at the two ATPase sites in all structures. This demonstrates of the high plasticity of the SMC1A HD/SMC3 HD interface and the capacity of the ATP gate to remain shut despite large internal reorganizations of the ATPase module.

Comparison of our engaged and open-engaged complexes revealed major differences. Notably, both lever effects are lost in the open-engaged complex. Consequently, the SMC1A and SMC3 second signature helices make large backward movements in the open-engaged complex compared to the engaged complex (Figure 6B-C). These movements are due to the back sliding of the SMC1A and SMC3 Helical-lobes with respect to their RecA-lobes. In the case of SMC1A, the resulting open-engaged SMC1A HD/RAD21C complex adopts a structure similar to our crystallographic structure of the independent SMC1A HD/RAD21C complex in its relaxed state when bound to ATPγS (Figure 6D). In this conformation, the rotational movement of the SMC1A CC from its central position, observed both in the apo and the engaged/clamped structures of the SMC1A HD, toward the RecA-lobe moves this CC towards the ATPase module DNA binding chamber.

In the case of SMC3, the open-engaged SMC3 GD also adopts a conformation similar to that of the SMC3 GD in the resting state (Figure 6E). However, the open-engaged SMC3 CC does not adopt a resting conformation, although this could be made possible due to the absence of a lever effect. Rather, upon loss of RAD21N, the SMC3 CC emerging from the SMC3 GD in the open-engaged state stiffens and makes a ∼90° angle compared to that in the resting state (Figure 6E). In this conformation, the open-engaged SMC3 CC is also repositioned toward the ATPase module DNA binding chamber (Figure 6E). As such, the movements of both SMC1A and SMC3 proximal coiled coils in the open- engaged state constricts the ATPase module DNA binding chamber (Figure 6F).

Despite these major structural rearrangements within both HDs, engagement is maintained in the absence of RAD21N. In addition, specific features observed upon engagement, such as the conformational rearrangement of the SMC3 wedge region and R-loop, are kept in the open-engaged ATPase module. However, in order to keep engagement, a major reorganization of the SMC1A HD/SMC3 HD interface occurs that is even larger than the one observed between the engaged and clamped ATPase modules (Figure 6G). This reorganization appears to further offset the ATP binding/hydrolysis motifs with respect to each other.

### The SMC3 HD/RAD21N interface including its RAD21N HelD domain are functionally important in vertebrates

Our different results have suggested an important role for the full N-terminal domain of RAD21 during the human Cohesin ATPase cycle. While the functional importance of the long RAD21N α-helix that binds to the SMC3 CC and enables the formation of the SMC3 HD/RAD21 interface has already been demonstrated, less is known on the functional importance of the RAD21N HelD domain. Sequence comparison shows that this domain is highly conserved in vertebrates (99% sequence identity) (Supplementary Figure 1C), suggesting that the role of the RAD21N HelD is conserved in these organisms.

Therefore, to assess the functional importance of the RAD21N HelD region in vertebrates, we obtained a characterized rad21a zebrafish mutant line (Xu et al., 2015) which we used to evaluate the effect of truncating various parts of the RAD21N region. Homozygous mutant larvae exhibited microcephaly, large pericardial edema and curved bodies at 3 days post-fertilization (dpf), which mirrored clinical features of the Cornelia de Lange Syndrome (CdLS) (Figures 7A and Supplementary Figure 8) (Kline et al., 2018). These phenotypes were specific and could be rescued efficiently upon injection of 200 pg of WT zebrafish full-length rad21a mRNA (Figures 7A-C and Supplementary Figure 8).

**Figure 7.**
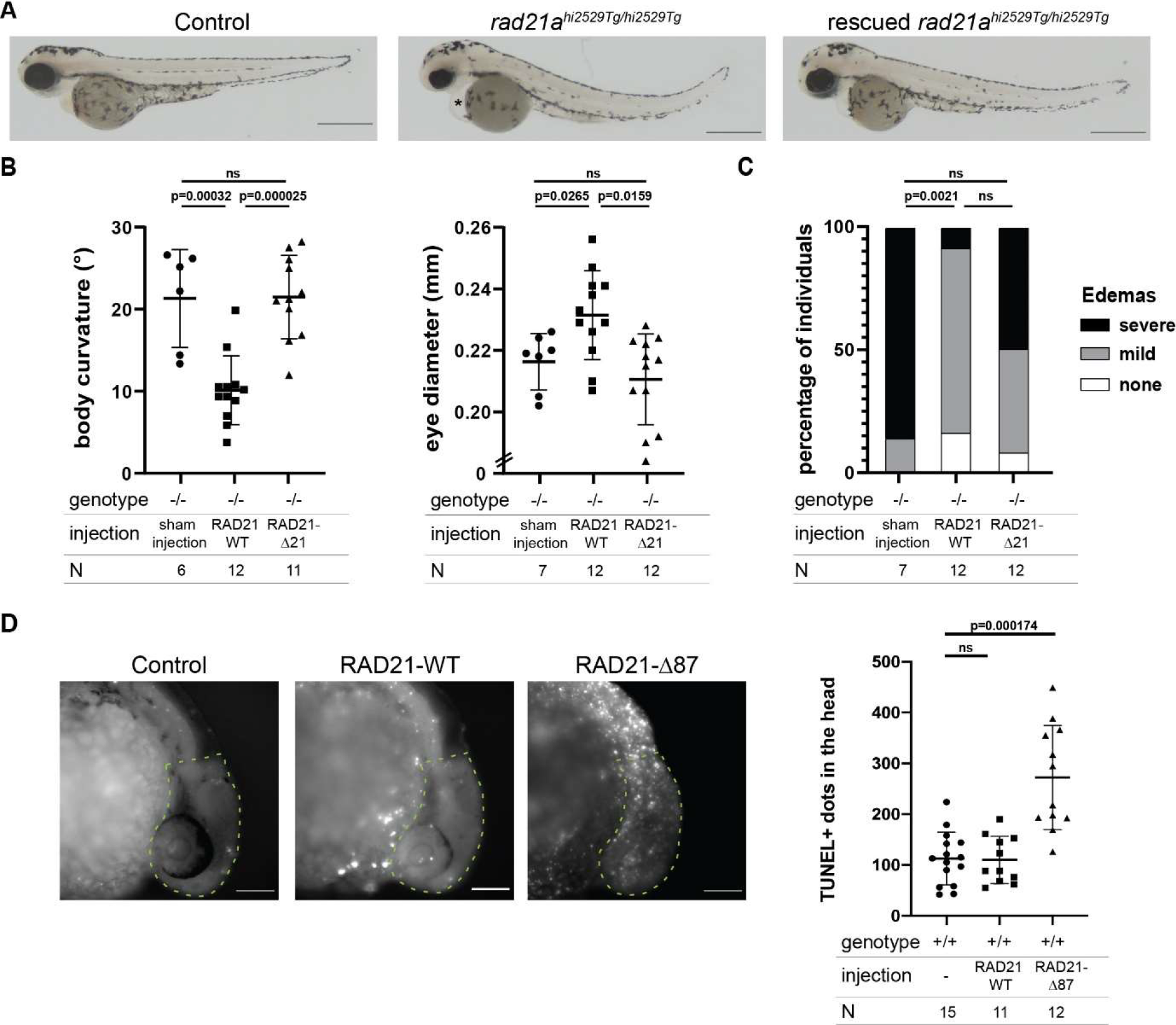
Functional importance of the RAD21N HelD domain and of the SMC3 HD/RAD21N interface in the zebrafish. A. From left to right, representative lateral images of zebrafish control rad21a+/+ (wildtype; WT), homozygous mutant rad21a^hi2529Tg/hi2529Tg^, and homozygous mutant rad21a^hi2529Tg/hi2529Tg^ larvae rescued with 200 pg of mRNA encoding WT full-length *Danio rerio* rad21a at 3 days post-fertilization (dpf). Scale bar, 0.5 mm. * indicates the presence of a pericardial edema. **B.** Dot plots measurements of body curvature (°) and eye diameter (mm) for homozygous mutant rad21a^hi2529Tg/hi2529Tg^ larvae injected with sham or full-length WT *Danio rerio* rad21a (RAD21 WT) or with the rad21a mRNA lacking the first 21 residues (RAD21-Δ21). Statistical significance was assessed by ANOVA followed by a Tukey’s test for post-hoc analysis. All experiments have been performed in biological duplicates. The p-values are indicated on the graphs. N corresponds to the number of embryos per condition. **C.** Bar graph showing the presence of pericardial edema for homozygous mutant rad21a^hi2529Tg/hi2529Tg^ larvae injected with sham or full-length WT *Danio rerio* rad21a (RAD21 WT) or with the rad21a mRNA lacking the first 21 residues (RAD21-Δ21). Larvae were binned into three categories: severe, mild or absent (none). Statistical significance was assessed by a Fisher’s exact test. All experiments have been performed in biological duplicates. The p-values are indicated on the graphs. N corresponds to the number of embryos per condition. **D.** From left to right, representative lateral images of uninjected wild-type (WT) larvae (shown as Control) and WT larvae injected with full length WT *Danio rerio* rad21a (RAD21 WT) or with the rad21a mRNA lacking the first 87 residues (RAD21-Δ87) at 1 dpf. The dot plot represents the number of TUNEL positive dots in the highlighted brain area (dotted line on the images) for the three conditions. A t-test was conducted between pairs of conditions to determine significance. The p-values are indicated on the graphs. The TUNEL assay has been repeated twice. N corresponds to the number of embryos per condition. Scale bar, 0.1 mm. ns: non-significant.

To test the requirement of the RAD21 HelD domain in Cohesin function, and notably of its N-terminal helix that enables the resting state conformation of the SMC3 HD, we injected a mutant mRNA that did not code for the first 21 amino acids of RAD21 (RAD21-Δ21) into homozygous mutant eggs. The RAD21-Δ21 mRNA failed to rescue the RAD21-/- phenotypes compared to the wild-type mRNA (Figures 7B-C and Supplementary Figure 8). Specifically, body curvature, eye diameter (as a read-out for microcephaly) and pericardial edema of RAD21-Δ21-injected larvae were affected indistinguishably from non-injected rad21a homozygous mutant larvae, suggesting that RAD21-Δ21 acts as a loss-of-function allele. The loss of rescue observed with the RAD21-Δ21 mRNA indicated that the absence of the first 21 residues induces a loss of function of rad21a, thus confirming the critical role of the RAD21 HelD domain for Cohesin function in vertebrates.

We made a comparative analysis with a rad21a mRNA lacking the full RAD21N region (RAD21-Δ87). This mutant, which should prevent the formation of the Cohesin complex *in vivo* by destroying the SMC3 HD/RAD21N interface, led to an aggravated phenotype rather than a rescued phenotype, as shown by the gross observation of dead tissue in the head of homozygous mutant embryos at 1 dpf, and developmental arrest. Similarly, injection of the zebrafish RAD21-Δ87 mRNA into wild-type eggs led to the same severe phenotype. The presence of dead cells was confirmed by acridine orange staining of RAD21-Δ87 mRNA-injected embryos (Supplementary Figure 8E). Accordingly, TUNEL staining revealed a significant increase of the number of apoptotic cells in the head compared to control embryos or embryos injected with WT rad21a mRNA (Figure 7D). These *in vivo* data suggest the essential function of RAD21N in vertebrates, including its N-terminal HelD domain.

## Discussion

### The SMC1A and SMC3 ATPase head domains undergo specific but concerted conformational changes during the Cohesin ATPase cycle

Our understanding of the molecular mechanisms underlying the Cohesin ATPase cycle has remained scarce although this cycle supports most Cohesin mechanisms and functions. The required flexibility of the Cohesin renders difficult the structural characterization at high resolution of this complex. Accordingly, cryo-electron microscopy studies using the full-length Cohesin complex have only provided high resolution structural data for the ATPase module bound to NIPBL^Scc2^ and DNA, revealing at the same time the strong flexibility of the regions linking the SMC1A and SMC3 HDs to the rest of the Cohesin complex (Collier et al., 2020; Higashi et al., 2020; Petela et al., 2021; Shi et al., 2020).

Here, we have studied the conformational dynamics of the Cohesin SMC1A and SMC3 ATPase head domains (HDs) along this cycle. Our results show that these HDs can recapitulate the ATPase activity of the full-length Cohesin core complex and provide high resolution structural data on major stages of the Cohesin ATPase cycle, including on the engaged ATPase module that is central to this cycle. Our results further reveal how both HDs behave differently but in a concerted manner during this cycle. Specifically, while both HDs have retained some similar mechanisms, most of our findings unravel mechanisms that are specific to each HD, which agrees with their evolutionary divergence at the sequence, structural and functional levels.

Differences are already observed at the nucleotide binding stage with different thermodynamic and regulatory mechanisms. At the structural level, while the resting conformation of the SMC3 HD is not significantly affected by nucleotide binding, the relaxed SMC1A HD undergoes specific rotational movements of its Helical-lobe with respect to its RecA-lobe. Upon engagement, a lever effect is observed in both HDs that causes a repositioning of both CCs. However, the SMC1A CC remains conformationally stable whereas the SMC3 CC/RAD21N complex oscillates. Nevertheless, the conjoint movements of the SMC1A and SMC3 CC upon engagement are concerted to form the engaged Cohesin ATPase module that can accommodate an incoming DNA molecule.

The oscillation of the SMC3 CC/RAD21N complex upon engagement builds on the intrinsic flexibility of the SMC3 CC that also enables the formation of the resting state of the independent SMC3 HD/RAD21N complex. Unexpectedly, the conformational flexibility of the SMC3 CC is lost upon dissociation of RAD21N from the ATPase module. In this case also the conjoint movements of the SMC1A and SMC3 CCs are concerted to form the open-engaged complex. In this case, however, these movements constrict the DNA binding chamber.

In the open-engaged state, both HDs return to conformations observed prior to engagement, still maintaining their ATP-dependent interaction. Upon ATP hydrolysis, the SMC1A and SMC3 HDs also return to conformations observed prior to engagement but cannot maintain engagement in the presence of ADP. This highlights the importance of the ATP γ-phosphate to keep the heterodimeric engagement of both HDs. This is, however, only rendered possible by the high plasticity of the SMC1A HD/SMC3 HD interface that enables the HDs to remain engaged while undergoing major structural rearrangements. The modular nature of both ATP binding sites, which are formed by small motifs whose positioning relative to each other can be varied, appears essential to allow this plasticity of the SMC1A HD/SMC3 HD interface, while also potentially regulating the Cohesin ATPase activity.

### The engaged Cohesin ATPase module needs to be stabilized to reach the clamped state

The structures of the yeast and human clamped complexes have shown how NIPBL^Scc2^ tightly clamps the DNA together with the ATP-engaged Cohesin ATPase module (Collier et al., 2020; Higashi et al., 2020; Shi et al., 2020). Our structures of the Cohesin ATPase module reveal, however, that binding of NIPBL^Scc2^ to the ATPase module to form the clamped complex requires specific structural rearrangements within this module. Specifically, while the ATP-dependent formation of the ATPase module already induces significant changes that appear important for NIPBL^Scc2^ docking, those are not sufficient for full NIPBL^Scc2^ binding to both SMC1A and SMC3 GDs and CCs.

Notably, we show that engagement specifically repositions and restructures the SMC1A F-loop that represents a crucial binding site of NIPBL^Scc2^. These changes are not only important for the proper docking of NIPBL^Scc2^ onto the ATPase module. They also ensure that NIPBL^Scc2^ will be correctly positioned for interacting through its E-loop with the SMC3 GD and through the C-terminal part of its hook with the SMC1A CC (Shi et al., 2020). Moreover, the docking of NIPBL^Scc2^ at the F-loop also appears to cause a reorganization of the SMC1A HD/SMC3 HD interface that also affects the positioning with respect to each other of the composite ATP binding and hydrolysis motifs at both ATPase sites.

The oscillation of the SMC3 CC/RAD21N complex upon engagement appears more difficult to overcome to reach the interaction observed between the SMC3 CC/Joint element and the NIPBL^Scc2^ nose in the clamped state. Notably, any intermediate conformation between the resting and clamped conformation of the SMC3 CC/RAD21N complex is sterically incompatible with the position of the NIPBL^Scc2^ nose in the clamped state (Supplementary Figure 9A). This suggests that the formation of the clamped complex requires a conjoint movement of the SMC3 CC/RAD21N complex and of the nose of NIPBL^Scc2^. Actually, the DNA in the clamped conformation plays a crucial role in positioning the NIPBL^Scc2^ nose for correct interaction with the SMC3 CC/Joint element.

Alternatively, a swing-and-clamp mechanism has been proposed for forming the clamped complex (Bauer et al., 2021). In this case, NIPBL^Scc2^ bound to DNA is proposed to first interacts with the SMC3 HD prior to the ATP-dependent engagement of both HDs. This mechanism is also compatible with our data. Notably, the resting SMC3 HD/RAD21N complex displays a positively charged electrostatic patch different from that observed in the clamped state (Supplementary Figure 9B-C) and that could serve as specific docking platform for the incoming NIPBL^Scc2^/DNA complex. The interactions of NIPBL^Scc2^ with SMC1A then occur in a second step following engagement. The importance of our observed engagement-induced reorganization of SMC1A remains valid in this mechanism. Moreover, the engagement-induced oscillation of the SMC3 CC/RAD21N complex should facilitate the repositioning of the SMC3-bound NIPBL^Scc2^ to interact with the SMC1A HD.

Following ATP hydrolysis, the concerted return of the SMC1A and SMC3 HDs to their relaxed and resting conformations is expected to release the tight grip on the DNA by opening the ATPase gate, but could also cause the disengagement of NIPBL^Scc2^ by affecting its docking sites, notably the SMC1A F-loop. It cannot be excluded, however, that NIPBL^Scc2^ could remain bound to the SMC1A HD by preserving the engaged conformation of this HD. Interestingly, in the open-engaged state, the SMC1A HD also adopts a relaxed conformation and the SMC3 CC has a different conformation. This suggests that a stable binding of NIPBL^Scc2^ to the ATPase module is less likely to occur in the open-engaged state, even if engagement is retained. Reassociation of RAD21N to the SMC3 CC could however induce the required changes that could lead to NIPBL^Scc2^ binding.

### The SMC1A and SMC3 HDs conformational flexibility serves to remodel the ATPase module DNA binding chamber

So far, binding of DNA by the Cohesin ATPase module has only been characterized structurally in the clamped state, revealing a DNA binding chamber formed by the GDs and proximal CCs of the engaged SMC1A and SMC3 HDs (Collier et al., 2020; Higashi et al., 2020; Shi et al., 2020). Specifically, the DNA lays on the positively charged pit formed by the engaged SMC1A and SMC3 GDs (Supplementary Figure 9D). In addition, the DNA tightly interacts with the SMC3 CC and the HelD of RAD21N. Notably, this latter domain displays a highly positive charge and appears to contribute strongly to the electrostatic binding and curvature of the DNA (Supplementary Figure 9D). In contrast, the SMC1A CC shows no strong electrostatic potential and appears mainly to act as a rigid wall to keep the DNA in its binding chamber.

In the engaged state, the oscillation of the SMC3 CC/RAD21N complex causes a widening of the DNA binding chamber (Supplementary Figure 9E). This potentially provides a way to facilitate DNA entry into this chamber, the electrostatic properties of the charged pit formed by both GDs and of the SMC1A CC remaining similar. However, this oscillation of the SMC3 CC/RAD21N complex also displaces the HelD and its basic patch that cannot bind tightly to the DNA (Supplementary Figure 9E). The electrostatic complementarity between the HelD and the DNA could lead to a stabilization of the SMC3 CC as observed in the clamped state. Our crosslinking experiments show, however, that such a stabilization by the DNA alone is most likely weak, further highlighting the role of NIPBL^Scc2^ in tightly keeping the DNA into its binding chamber. In addition, as shown by our crosslinking experiment, the amplitude of the oscillation of the SMC3 CC/RAD21N complex is most likely small and might even be more restricted in the case of the full Cohesin core complex. This oscillation should however prevent the tight clamping of the DNA until the clamped complex is formed.

Although opening of the SMC3 HD/RAD21N interface stabilizes the SMC3 CC, it causes its positioning, as well as that of SMC1A, inside the DNA binding chamber. The resulting constriction of this chamber is poorly compatible with DNA binding. Moreover, the electrostatic potential of the DNA binding chamber in the open-engaged complex displays a reduced positive charge (Supplementary Figure 9F) and the access to this chamber is restricted due to the closed ATP gate. Interestingly, fitting of our human open-engaged structure within the cryo-EM map of the full-length ATP-engaged *S. cerevisiae* core Cohesin complex lacking Scc1 (EMBD entry EMD-12889) reveals that the conformation of the open-engaged SMC3 CC agrees relatively well with the cryo-EM density (Supplementary Figure 10). Specifically, this yeast complex forms a compact and overall rigid complex compared to other structures published (Collier et al., 2020; Higashi et al., 2020; Petela et al., 2021; Shi et al., 2020).

The SMC3 HD/RAD21N interface has been termed DNA exit gate and its opening has been shown to allow the release from the Cohesin ring of topologically entrapped DNA (Beckouet et al., 2016; Buheitel and Stemmann, 2013; Chan et al., 2012; Eichinger et al., 2013; Huis in ’t Veld et al., 2014; Murayama and Uhlmann, 2015). In addition, entrapment of DNA has been shown to occur in the kleisin compartment but not in the SMC compartment (Chapard et al., 2019). Opening of the SMC3 HD/RAD21N interface could therefore serve two objectives. The first objective would be to release DNA. The second objective could be to force the SMC heterodimer to adopt a tight conformation less prone to interactions with DNA, to prevent DNA interaction and potentially entering in the SMC compartment.

### Importance of RAD21N and its HelD domain

Our study has highlighted the role played by RAD21N, including its HelD domain, for the specific conformational dynamics of the SMC3 HD and Cohesin ATPase module. Our *in vivo* experiments in the zebrafish confirm that RAD21N and its HelD domain are functionally important, including the first 21 residues of RAD21 that interact with the SMC3 GD in the resting conformation of the SMC3 HD/RAD21N complex (Supplementary Figure 11A). Previous work has also already suggested a role of the first 20 amino acids of RAD21 in the interplay of this kleisin with PDS5 and WAPL (Ouyang et al., 2016). In addition, the HelD domain is also involved in the formation of the clamped state, both in human and in *S. pombe* (Supplementary Figure 11B-C) (Higashi et al., 2020; Shi et al., 2020). Curiously, the HelD domain is less structured in *S. cerevisiae* (Supplementary Figure 11D-E) (Collier et al., 2020; Gligoris et al., 2014). This does not, however, prevent the formation of the *S. cerevisiae* clamped state. The role of the HelD domain appears therefore dedicated to specific mechanisms that remain to be further characterized.

In the paralogous Condensin complex, the CAP-H^Brn1^ kleisin also harbours a HelD domain that shows sequence and structural similarities with the RAD21^Scc1^ HelD, including a N-terminal α-helix and a lysine equivalent to K26 (CAP-H^Brn1^ K42; Supplementary Figure 11F-G). In addition, the structures of the yeast Condensin ATPase module bound to DNA and to the regulatory subunit Ycs4 show significant similarities with the Cohesin clamped structures, including the interaction of the CAP-H^Brn1^ HelD with the DNA (Lee et al., 2022; Shaltiel et al., 2022). It is therefore likely that the mechanisms underlying the ATPase cycles of Cohesin and Condensin share strong similarities. Specifically, a flexibility of the SMC2 CC at a similar position than that observed for the SMC3 CC has also been observed (Hassler et al., 2019; Lee et al., 2020).

Collectively, our results reveal the specific structural movements and conformational rearrangements undergone by the SMC1A and SMC3 HDs upon ATP binding, ATP-dependent engagement, ATP hydrolysis and opening of the SMC3 CC/RAD21N interface nd how these are concerted. Our result also suggest that the ATP gate can potentially be kept closed in a low ATPase activity conformation through specific regulatory mechanisms. In this respect, the Cohesin ATPase cycle can be divided in two potentially independent steps: an ATP-dependent engagement with formation of the ATP gate and an ATP hydrolysis-dependent disengagement causing the opening of the ATP gate. The specific movements and rearrangements we describe should impact all Cohesin mechanisms that rely on the Cohesin ATPase cycle.

## Acknowledgments

We thank Catherine Birck, Alastair Mc Ewen, Nils Marechal and Pierre Poussin-Courmontagne of the IGBMC Integrated Structural Biology platform for their help during biochemical, biophysical and structural data collection and analysis. We thank Gabor Papai, Nils Marechal and Albert Weixlbaumer for advice on cryo-EM structure refinement. We thank Piotr Sosnowski and Helgo Schmidt for advice with the ATPase assays. We thank the Zebrafish International Resource Center (ZIRC, Oregon, US) for providing the zebrafish line used in this study, and the IGBMC Zebrafish Facility, in particular Sandrine Geschier, for maintenance and care of the zebrafish line. We thank Gaëlle Hayot for help with statistical analyses. We thank members of the SOLEIL and SLS synchrotrons for the use of their beamline facilities and for help during data collection. This work was supported by the Agence Nationale de la Recherche (grant numbers ANR-10-LABX-030-INRT to M.V.G.; ANR-17-EURE-0023 to M.V.G.; ANR-10-IDEX-0002 to M.L.D.D. and to C.R.; ANR-17-CE12-0006 to C.G.; ANR-10-INBS-0005-01), the Fondation Association pour la Recherche sur le Cancer (grant numbers ARCPJA20181208268 to C.R.; ARCPJA2021060003715 to C.R.; DOC20180507150 to P.L.), the Fondation pour la Recherche Médicale (grant number FDT202106012973 to P.L.), the INSTRUCT-ERIC, the European Regional Development Fund, the Alsace Region, the General Council of Bas-Rhin, the French Ministry of Higher Education and Research, and by institutional funds from the Centre National de la Recherche Scientifique (CNRS), the Institut National de la Santé et de la Recherche Médicale (INSERM) and the University of Strasbourg.

## Methods

### Cloning

The SMC1A and SMC3 ATPase head domain (HD) constructs were generated by PCR using the full length human *smc1a* and *smc3* genes as templates. The sequences coding for the hinges and for the coiled-coils up to the joint elements were replaced by sequences coding for a short protein linker (either ESSKHPASLVPRGS or GSGSLVPRGSGS), as previously reported (Gligoris et al., 2014; Haering et al., 2004). The RAD21N and RAD21C constructs were generated by PCR using the full length human *rad21* gene as template. Point mutations were introduced into the constructions using rolling circle or nested PCR strategies. The constructs were cloned by Gibson assembly into bacterial co-expression vectors (Diebold et al., 2011; Fribourg et al., 2001; Romier et al., 2006; Vincentelli and Romier, 2016) to code either for native SMC1A HD (SMC1ACCsh, SMC1ACC, SMC1AJ constructs) and SMC3 HD (SMC3CC and SMC3J constructs) or for RAD21 constructs (RAD21N and RAD21C) followed by a 3C protease cleavage site and a 10xhistidine fusion tag.

### Protein complexes production and purification

The same large-scale overproduction and purification methods have been used for all protein complexes, unless stated. The SMC1A or SMC3 HDs-coding plasmids were respectively co-transformed with the RAD21C- or RAD21N-coding plasmids into chemically competent Escherichia coli BL21(DE3) cells (Novagen). Co-transformed cells were selected using the appropriate antibiotics. Colonies were used to inoculate large cultures of 2x LB medium that were grown for 6 hours at 37°C. Protein expression was induced at 25°C by the addition of a final concentration of 0.7 mM Isopropyl β-D-1- thiogalactopyranoside (IPTG; Euromedex), and cells were further grown overnight at 25°C. Culture media was discarded after centrifugation and the bacterial pellets from 3 L of culture were resuspended with 30 ml lysis buffer containing either 200 mM or 500 mM NaCl and 10 mM Tris-HCl pH 8. Pellets were stored at -20°C until further use.

After sonication, lysate clarification was performed by centrifugation (1 hour at 17,000 rpm). The recombinant SMC1A HD/RAD21C and SMC3 HD/RAD21N protein complexes were then purified by affinity chromatography using the 10xHis purification tag on RAD21 by incubating the cleared lysates with TALON Metal Affinity Resin (Takara Bio). The purification tag was then removed on the affinity beads by overnight 3C protease digestion at 4°C. Removal of nucleic acid contaminants was performed using 1 ml or 5 ml HiTrap Heparin column (GE Healthcare) and eluted using a NaCl gradient from 50 mM to 1 M. Peak fractions containing the protein complexes were pooled and further purified by size exclusion chromatography using a 16/60 Superdex 200 column (GE Healthcare) equilibrated with a buffer containing 200 mM NaCl, 10 mM Tris-HCl pH 8.0, 2 mM MgCl2 and either 1 mM TCEP (samples for crystallization and ITC assays) or 2 mM DTT (samples for ATPase assays). The main peak fractions with the protein complexes were pooled, concentrated with AMICON Ultra concentrator filters (Merck Milipore), and either used immediately or frozen in liquid nitrogen and stored at -80°C for later use.

### ATPase assays

Measurement of the ATPase activity of wildtype and mutant SMC1A-HD/RAD21C and SMC3- HD/RAD21N complex were assayed using the EnzChek Phosphate Assay Kit (Thermo Fischer Scientific). For these assays, each protein sample (final concentration of 10 µM) was incubated with 1 mM ATP. After addition of ATP, the ATP hydrolysis activity was immediately assessed at 30°C, by measuring the absorbance at 360 nm every 42 seconds for 2 hours, using a spectrophotometer plate reader (TECAN). The ATPase activities, expressed in Pi molecules released per dimer and per minute shown in the figures were calculated in the linear range of the curves. These experiments have been performed in triplicates.

### Isothermal titration calorimetry measurements

For ITC measurements, ATP, ATPγS or ADP were dissolved at a final concentration of 4 mM into the protein gel filtration buffer (200 mM NaCl, 10 mM Tris-HCl pH 8, 2 mM MgCl2 and 1 mM TCEP). Each nucleotide was injected into 189 to 330 µM of either the wildtype or the EQ-mutant SMC1ACC/RAD21C or SMC3CC/RAD21N complexes, respectively. ITC measurements were performed at 5 °C using a PEAQ-ITC microcalorimeter (Malvern Panalytical). ITC data were then corrected for the dilution heat generated by the injection of the buffer into the protein sample and of the nucleotide sample into the buffer.

ITC data were then fitted using the AFFINImeter analysis software. In most measurements, a single binding event was observed and processing was performed for a single binding event, which gave a good fitting of the experimental data. In the case of the SMC3CC/RAD21N complex two events were observed and the processing was made with a two binding events model. Since ADP binding to this complex led to a single event, we reasoned that the second binding event could be due to a homodimerization at high concentration of the SMC3CC/RAD21N complex in the presence of ATP and ATPγS. We therefore processed the ITC data for SMC3 with a model including two consecutive binding events: an initial ATP-binding event followed by a homodimerization event. This model gave a good fitting of the experimental data and yielded one Kd in the µM range and another in the low nM range for the two events. Since the measured Kd of ADP for SMC3CC/RAD21N was in the µM range, we assumed that the Kd of ATP and ATPγS for the same complex also corresponded to the higher measured Kd. All measured thermodynamic parameters are given in Supplementary Table 1

### Cross-linking experiments

For crosslinking experiments, all endogenous cysteines of the SMC1ACC, SMC3CC, RAD21N and RAD21C constructs were mutated into serines to avoid initially observed unspecific crosslinking. The specific cysteines were then introduced in the SMC3CC (D92C and D120C) and RAD21N (K25C or K26C) constructs. All mutated complexes were purified following standard protocol described above. Mutated SMC3CC/RAD21N complexes were mixed to a final concentration of 6 µM alone or in the presence of SMC1ACC/RAD21C (7 µM), ATP (20 µM), DNA (10 µM) and MgCl2 (40 µM). Reactions were incubated for 5 min on ice or at room temperature before addition of 0.5 mM BMOE or DMSO. The reaction was quenched by addition of Laemmli buffer containing β-mercaptoethanol after 2, 5 and 10 minutes, and was further heated at 70°C for 5 min. The samples were loaded on acrylamide gel and band intensity was measured using Fiji (Schindelin et al., 2012). All experiments were performed in triplicates.

### Crystallizations

Initial crystallization assays were carried out with commercial crystallization screens in swissci 96-Well 3-Drop MRC crystallization plates (Molecular Dimensions). Crystals were grown using the sitting-drop vapor-diffusion method at 4, 20 and 27°C. Briefly, 200 nl of 4 to 15 mg/ml protein sample were mixed with 200 nl of reservoir solution. Several conditions from the commercial crystallization screens PACT, JCSG+, Classics, WIZARD I and II, BCS and LFS yielded protein crystals within a few hours up to several weeks. Some of the crystals, especially those from the SMC3 HD/RAD21N complex, required extensive optimization to improve their stability and diffraction limit. Selected crystals were cryo-protected with 20% (v/v) glycerol or 20% (v/v) PEG 200, then flash-cooled and stored into liquid nitrogen until data collection. Crystallization conditions are provided in Supplementary Tables 2, 3 and 4.

### Crystallographic structure determination, model building and refinement

X-ray diffraction data were collected at the SOLEIL and SLS synchrotrons. High resolution diffraction data (ranging from 1.4 to 3.1 Å) were obtained and processed by indexation, integration, and scaling within the XDS program (Kabsch, 2010). Data were merged using Aimless from the CCP4 software suite (1994). The various SMC1A HD/RAD21C and SMC3 HD/RAD21N structures were solved by molecular replacement using PhaserMR (McCoy et al., 2007) using respectively the yeast Smc1 HD/Scc1C (PDB entry 1W1W) and Smc3 HD/Scc1N (PDB entry 4UX3) structures as models. The initial models were subsequently iteratively built manually within Coot and refined using Phenix (Adams et al., 2010; Emsley et al., 2010). All refined models were verified with Molprobity (Williams et al., 2018) and showed good refinement statistics (Supplementary tables 2, 3 and 4).

### Reconstitution of the ATP-engaged ATPase module by size exclusion chromatography

For the characterization of the stably engaged ATPase module by size exclusion chromatography, the WT or EQ independently purified SMC1ACC/RAD21C and SMC3CC/RAD21N complexes were supplemented or not with 0.5/1 mM of the different nucleotides (ADP, ATP, ATPγS, AMP-PNP), diluted to a concentration of 50 µM, and loaded onto a Superdex S200 10/300 column (GE Healthcare) equilibrated with a buffer containing 200 mM NaCl, 10 mM Tris-HCl pH 8.5 mM MgCl2 and 1 mM TCEP, either without nucleotide or supplemented with 0.5/1 mM of the chosen nucleotide. The experiments performed with the WT and EQ, LV and DE mutants were performed initially using the same conditions but incorporating the SMC3J/RAD21N complex instead of the SMC3CC/RAD21N complex.

### Reconstitution of the ATP-engaged ATPase module by analytical ultracentrifugation (AUC) experiments

Analytical ultracentrifugation sedimentation velocity experiments were performed with a ProteomeLab® XL (Beckman Coulter) at 4°C and 42,000 rpm with absorbance detection at 280 nm. Independently purified SMC1ACC-EQ/RAD21C and SMC3CC-EQ/RAD21N complexes, stored in a buffer composed of 200 mM NaCl, 10 mM Tris-HCl pH 8.5 mM MgCl2 and 1 mM TCEP, were diluted to a final concentration of 1 mg/mL for experiments with the homodimers or mixed with a 1:0.9 ratio to a final concentration of 1 mg/mL for the heterodimers. ADP, ATP and low-hydrolysable ATP analogs (ATPγS, AMP-PNP) were added to the diluted samples at a concentration of 0.5/1 mM prior to measurements. The A280 nm scans data were acquired at 9 min intervals for 24h. The SEDNTERP software (Laue, 1992) was used to estimate the partial specific volume of the protein (υ), the density (ρ), and the viscosity (η) of the samples. Data were analyzed in the SEDFIT software (Schuck, 2000) using the continuous c(s) distribution analysis. AUC graphs were rendered in the GUSSI software (Brautigam, 2015).

### Cryo-electron microscopy sample preparation and data acquisition

For the preparation of the cryo-electron microscopy (cryo-EM) samples, the independently purified SMC1ACC-EQ/RAD21C and SMC3CC-EQ/RAD21N samples were mixed with a 1:0.9 ratio and loaded onto a Superdex 200 16/600 column (GE Healthcare) equilibrated with the purification buffer (200 mM NaCl, 10 mM Tris-HCl pH 8.2 mM MgCl2 and 1 mM TCEP) supplemented with 1 mM ATP. The engaged complex obtained from the most concentrated chromatographic fraction was diluted to 0.3 mg/mL. A 3 μl aliquot of the sample was applied onto C-flat, Gold or in-house PEGylated gold 1.2/1.3 grids (Meyerson et al., 2014). The grid was blotted for 4s (blot force 5) and flash-frozen in liquid ethane using Vitrobot Mark IV (FEI) at 4°C and 100% humidity. Micrographs were acquired on a Glacios Cryo-TEM operated at 200kV on a K2 Summit camera in counting mode. Automated data acquisition was carried out using the SerialEM software (Mastronarde, 2005) at a 45,000 magnification in a nanoprobe TEM mode, which yielded a pixel size of 0.901 Å. The defocus range was set from -0.8 to -2.0 or -3.0. 40 movies frames were recorded at a dose rate of 7.4 electrons per Å2 per second to total doses of 43.61 up to 63.85 e/Å2. A total of 11,313 micrographs were collected. Detailed collection data are provided in Supplementary Table 5.

### Cryo-electron microscopy data processing, model building and refinement

For all micrographs, motion-correction, dose-weighting and CTF estimation was performed in Cryosparc. Particles were picked in Cryosparc using first a blob picker job, followed by 2D classification. Templates were selected for template-based picking in Cryosparc, yielding a dataset of 7,098,916 particles. Subsequently, particles were extracted with a box size of 256 px and binned to 128 px. Two rounds of 2D classification were performed to clean up the dataset, and 1,037,793 particles were kept and used to generate 5 ab-initio maps. Three of these maps resembled to the engaged head model while the others two showed no particular features. Two rounds of heterogenous refinement (with a box size of 64 px) were performed on the 5 maps to further clean up the dataset by keeping the particles belonging to the three recognizable classes generated by ab initio reconstruction. The remaining 652,840 particles were reextracted with a box size of 256 px and further cleaned up by two additional rounds of heterogenous refinement. Eventually, the 174,054 and 144,776 particles corresponding respectively to the two best models were used in a non-uniformed refinement job. However, the map of the second model showed a significant level of noise due possibly to over-refinement. The corresponding particles and metadata were therefore exported from Cryosparc to Relion using the pyem conversion script. These were then imported in Relion3 and a new 3D refinement was performed followed by postprocessing. This resulting map had clearly improved details and no sign of over-refinement. The two cryo-EM maps issued from these different refinement steps have resolutions of 3.6 Å (first model) and 4.4 Å (second model).

Model building was performed using our crystal structures of the SMC1ACC/RAD21C, SMC1ACCsh/RAD21C and SMC3CC/RAD21N for docking into the 3.6 Å resolution cryo-EM map using UCSF chimera (Pettersen et al., 2004). After manual adjustments, real-space refinement of the model was performed in Phenix (Liebschner et al., 2019) with secondary structure restraints, global minimization, morphing, and simulated annealing. The model was then further improved by several cycles of manual building/adjustment in Coot (Emsley et al., 2010) and real space refinement in Phenix. Model validation was performed within Coot and with MolProbity (Prisant et al., 2020). This model was then used for docking into the cryo-EM map at 4.4 Å resolution and this second model was modified and refined using the same procedures as the first one, its lower resolution limiting modelling but enabling positioning of the secondary structure elements.

### Zebrafish mutant line, in vivo complementation experiments and whole-mount immunostaining

Zebrafish (*Danio rerio*) were raised and maintained as described (Westerfield, 2007). The mutant line rad21a^hi2529Tg/+^ (AB genetic background) was obtained from the International Resource Center for Zebrafish (ZIRC#ZDB-ALT-041006-8). The zebrafish line reproduces normally. The rad21a heterozygote mutant exhibits no phenotype. The rad21a homozygote mutants died at 4 days post-fertilization (dpf) from severe pericardial edema. For this study, viable heterozygous mutant adults were crossed and obtained larvae were individually scored and genotyped following the procedure previously described (Xu et al., 2015). All the experiments were at least duplicated, all phenotypes were scored blind to the genotypes, and all statistical analyses were performed using GraphPad Prism v8.0.2.263 (GraphPad Software, San Diego, CA) and R (R Core Team (2021). R: A language and environment for statistical computing. R Foundation for Statistical Computing, Vienna, Austria. URL https://www.R-project.org/). For wild-type (WT) and mutant rescue experiments, *Danio rerio* WT and truncated rad21a constructs were cloned into the pCS2 vector, sequenced and transcribed using the SP6 mMessage Machine kit (Invitrogen). 200 pg RNA (WT or mutants) were injected into mutant or WT zebrafish embryos at the 1- to 2-cell stage. Eye diameter, body curvature and pericardial edema were scored on injected larvae at 3 dpf fixed in 4% PFA and washed in PBS-Tween 0.1%. Body curvature and eye diameter were measured using Fiji and statistical differences were assessed using ANOVA followed by Tukey’s test for post-hoc analysis. For edema scoring, embryos at 3 dpf were classified into three groups: none, mild, and severe based on the size of the edema compared with an age-matched control group from the same clutch and a Fisher’s exact test was performed to determine significance.

For whole-mount TUNEL assay, apoptotic cell death in 1-day old zebrafish whole-mounts was detected according to a modification of the ApopTag® Fluorescein In Situ Apoptosis Detection Kit (Merck Millipore) protocol. Dechorionated embryos were fixed in 4% paraformaldehyde (PFA) at 4°C overnight and stored in 100% methanol at -20°C. After rehydration in PBS, embryos were digested with proteinase K (10µg/ml) in PBS for 5 minutes at room temperature and washed 3 times in sterile water for 3 minutes each. Embryos were post-fixed with 4% PFA for 20 minutes at room temperature followed by a 10-minute incubation in prechilled ethanol:acetic acid (2:1) at -20°C. Embryos were washed in PBS-T (PBS 1X, 0.1% Tween-20) for 5 minutes, 3 times at room temperature. Incubation in equilibration buffer and further steps were followed according to the manufacturer instructions. Z-stack image acquisition was performed using a MacroFluo ORCA Flash macroscope (Leica). TUNEL staining was quantified by counting positive cells/dots in zebrafish head (forebrain, midbrain, eyes). Counting was performed using Fiji by combining automatic counting (ITCN plugin) with manual counting. Statistical differences were assessed with a t-test with Welch’s correction when necessary. For acridine orange staining, one-day old embryos were dechorionated and incubated at 28°C for 30 min in E3 embryo medium supplemented with 2 µg/mL acridine orange. After extensive washing, embryos were anesthetized with tricaine and Z-stack imaging was performed with GFP green light excitation. Cell counting was performed with Fiji using the ITCN plugin. Statistical differences were assessed with a t-test. All animal experiments were carried out according to the guidelines of the Ethics Committee of IGBMC and ethical approval was obtained from the French Ministry of Higher Education and Research under the number APAFIS#15025-2018041616344504.

