## Supplemental information for "The Cohesin ATPase cycle is mediated by specific conformational dynamics and interface plasticity of SMC1A and SMC3 ATPase domains"

### SUPPLEMENTARY DATA

A

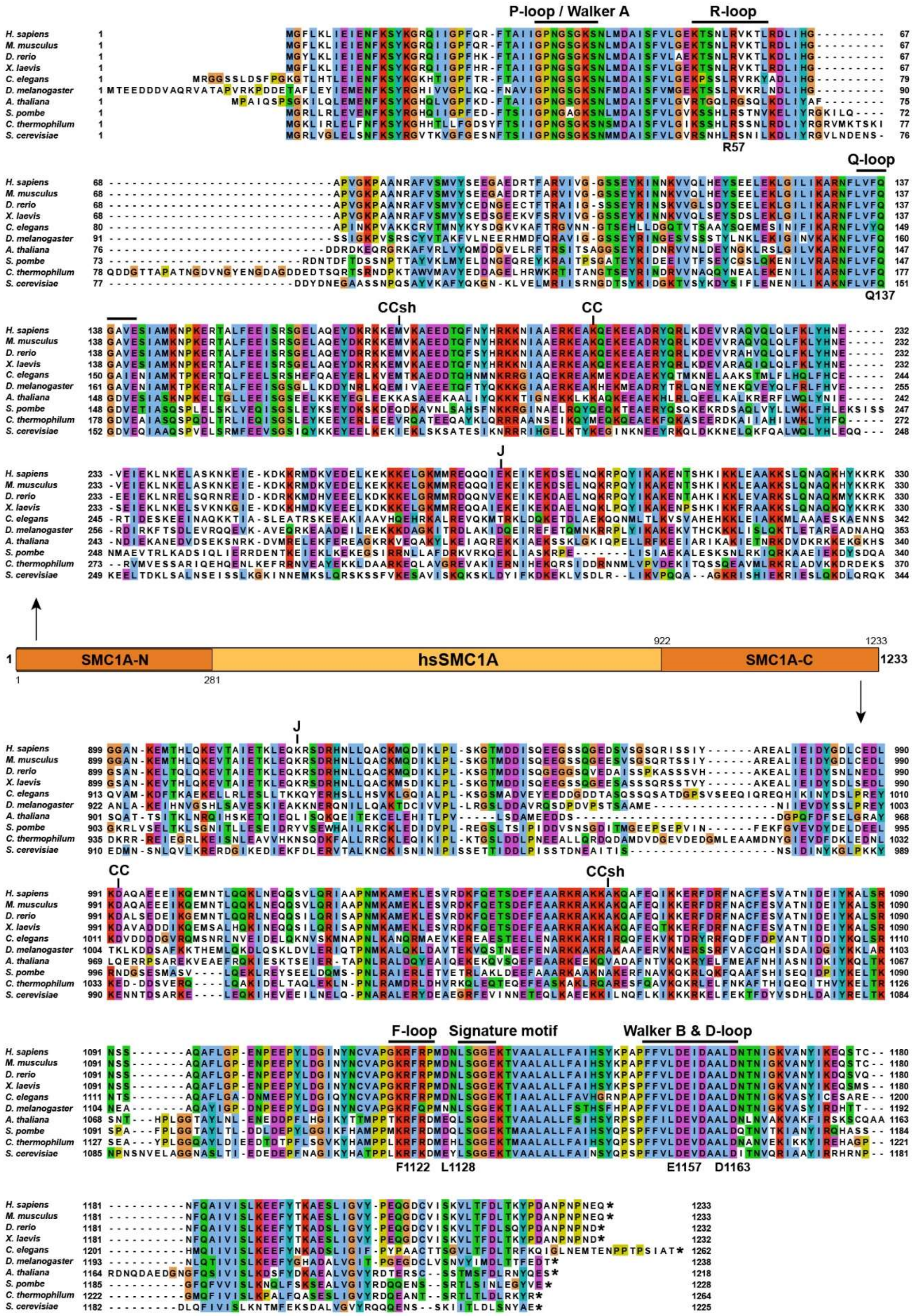

P-loop / Walker A
R-loop
R57
Q-loop
Q137
CCsh
CC
J
H. sapiens
M. musculus
D. rerio
X. laevis
C. elegans
D. melanogaster
A. thaliana
S. pombe
C. thermophilum
S. cerevisiae
SMC1A-N
hsSMC1A
SMC1A-C
1
281
922
1233
F-loop
Signature motif
Walker B & D-loop
F1122
L1128
E1157
D1163

B

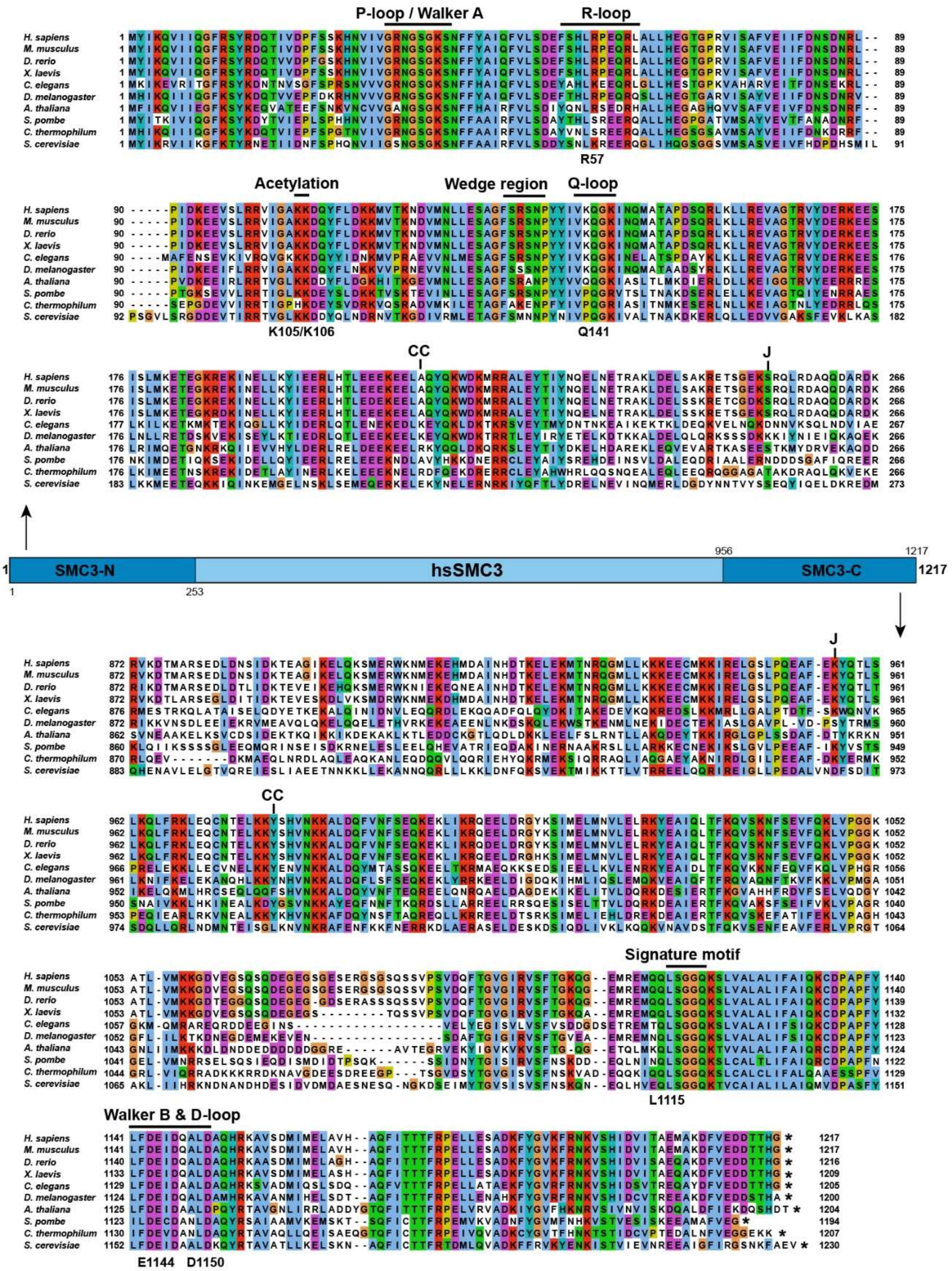

P-loop / Walker A
R-loop
R57
Acetylation
Wedge region
Q-loop
K105/K106
Q141
CC
J
SMC3-N
hsSMC3
SMC3-C
Signature motif
L1115
Walker B & D-loop
E1144
D1150
H. sapiens
M. musculus
D. rerio
X. laevis
C. elegans
D. melanogaster
A. thaliana
S. pombe
C. thermophilum
S. cerevisiae

**C**

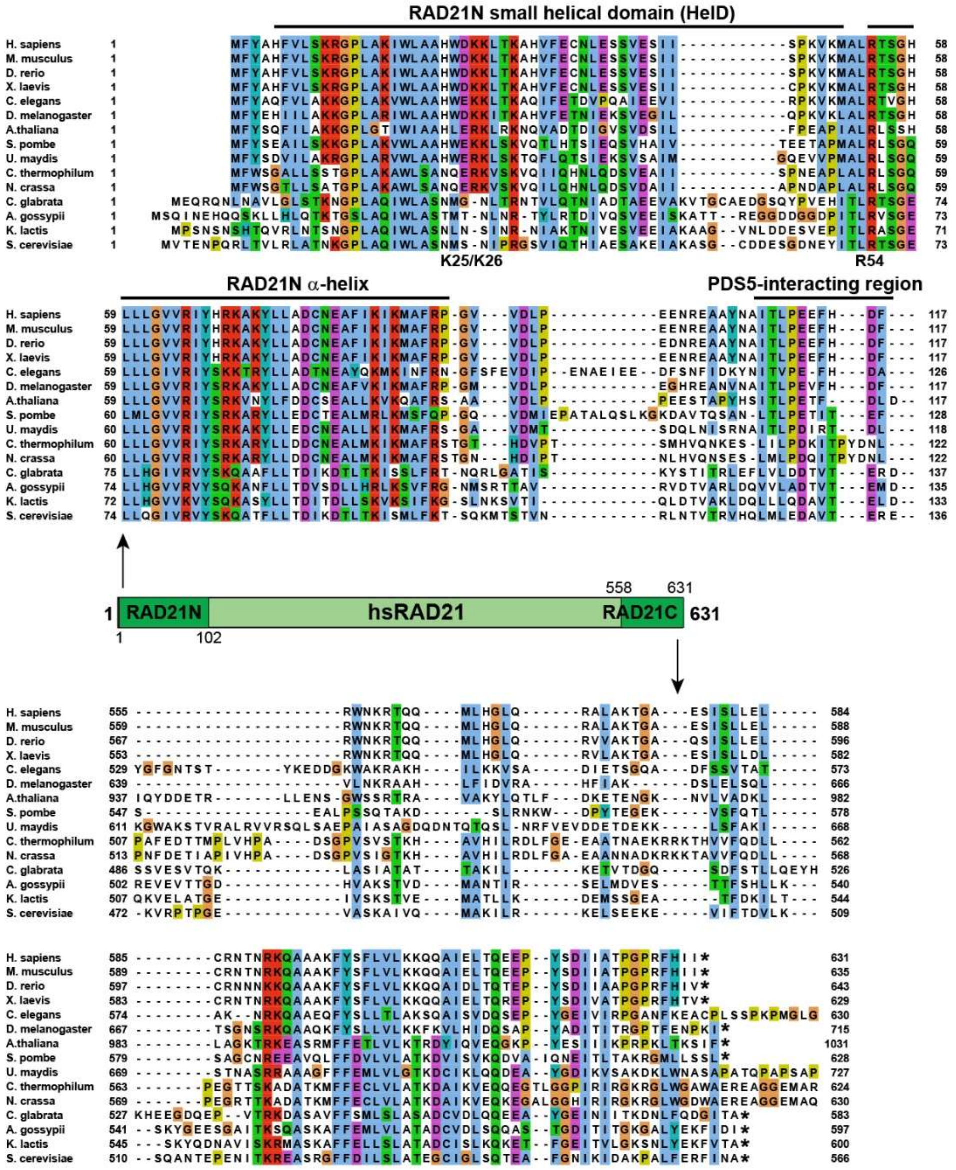


**Supplementary Figure 1. Sequence alignments of the SMC1A HD, SMC3 HD, RAD21N and RAD21C regions. A.** Sequence alignment of the N- and C-terminal semi-ATPase domains of SMC1A from various organisms. The clustalX coloring scheme is used for representing sequence conservations. The major sequence/structural elements and residues discussed in the text are indicated. Asterisks indicate the C-terminal end of the sequences. Boundaries of the constructs used in the study (CCsh, CC, J) are marked. In the central representation of the full-length SMC1A, the constructs used are represented at both extremities and the numbering of the longer construct is indicated. **B**. Same as in (A) for the N- and C-terminal semi-ATPase domains of SMC3. **C**. Same as in (A) for the N- and C-terminal domains of RAD21$^{Scc1}$.

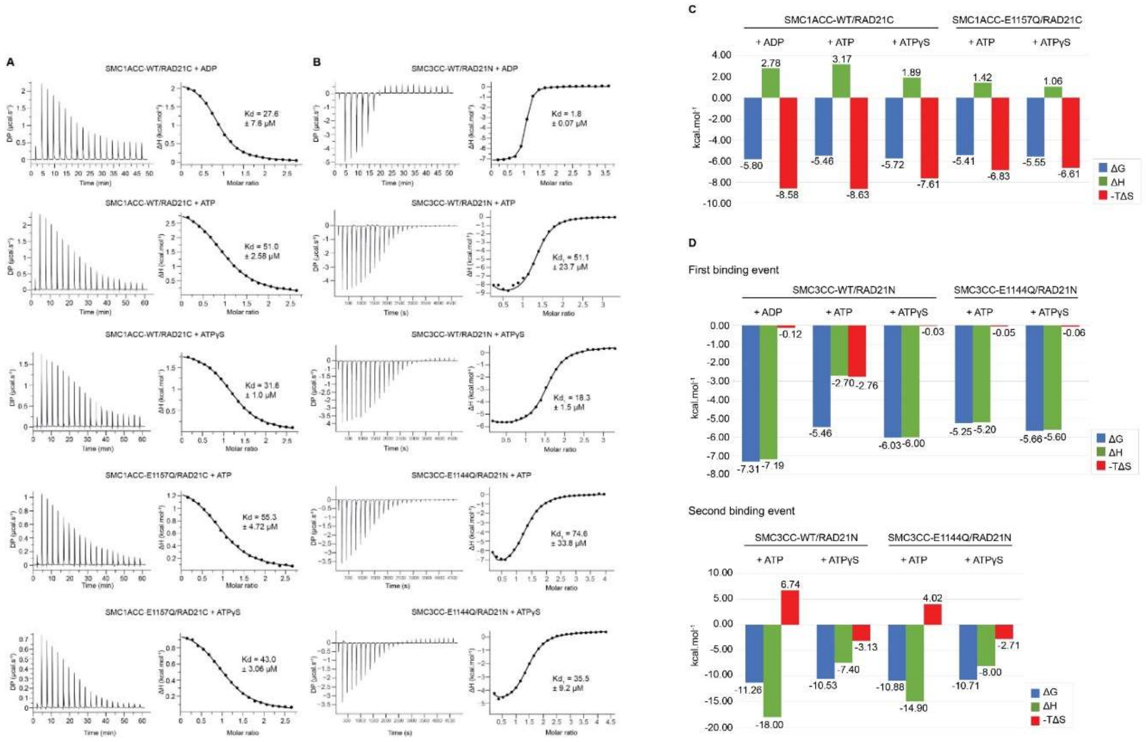


**Supplementary Figure 2. Analysis by isothermal titration calorimetry (ITC) of ADP, ATP and ATPγS binding to the SMC1A HD and SMC3 HD. A.** Measurements of the Kd of ADP, ATP and ATPγS for the SMC1ACC/RAD21C complex using the SMC1ACC WT and E1157Q mutant constructs. The measurements profiles are provided on the left panels, the interpolations on the right panels. The Kd values are given above the interpolation curves. The complete thermodynamic parameters obtained from these experiments are given in Supplementary Table 1. **B**. Same as in (A) for the SMC3CC/RAD21N complex with the WT and E1144Q mutant constructs. The complete thermodynamic parameters obtained from these experiments are given in Supplementary Table 1. **C**. Thermodynamic signatures obtained from the ITC experiments performed with the SMC1ACC/RAD21C complex. **D**. Same as in (C) for the SMC3CC/RAD21N complex.

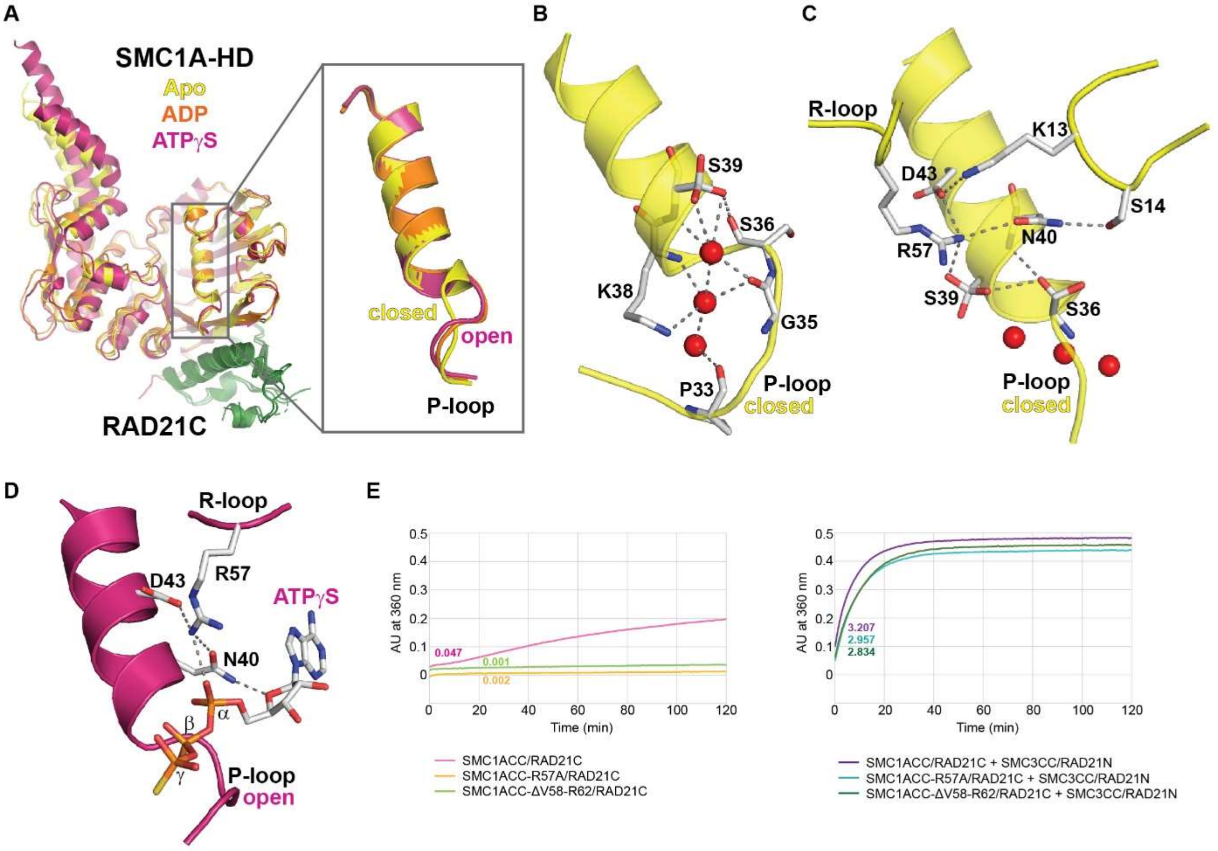


**Supplementary Figure 3. Organization of the SMC1A HD ATP binding site by the P-loop and R-loop.** **A.** Ribbon representation of the SMC1ACC/RAD21C complex in apo, ADP-bound and ATPγS-bound forms upon superposition of the α-helix following the P-loop. In the apo form, the P-loop adopts a closed conformation, whereas it has an open conformation in the nucleotide-bound forms. **B.** Interaction network, including some water molecules (red spheres), that stabilize the apo closed conformation of the P-loop. **C.** Interaction network that stabilizes R57 from the SMC1A R-loop and enables this residue to participate to the organization of the SMC1A ATP binding site prior to nucleotide binding. **D.** Interaction of SMC1A R57 with a bound ATPγS molecule that enables this residue to stabilize this nucleotide in its binding pocket. **E.** ATPase activity of the independent (left panel) and SMC3-bound (right panel) SMC1A R57A and CdLS ΔV58-R62 mutants. While the mutations hamper ATP hydrolysis by the independent SMC1A HD, they only slightly affect the ATPase activity of the engaged complex. The ATPase activity is given in Pi molecules released per dimer and per minute.

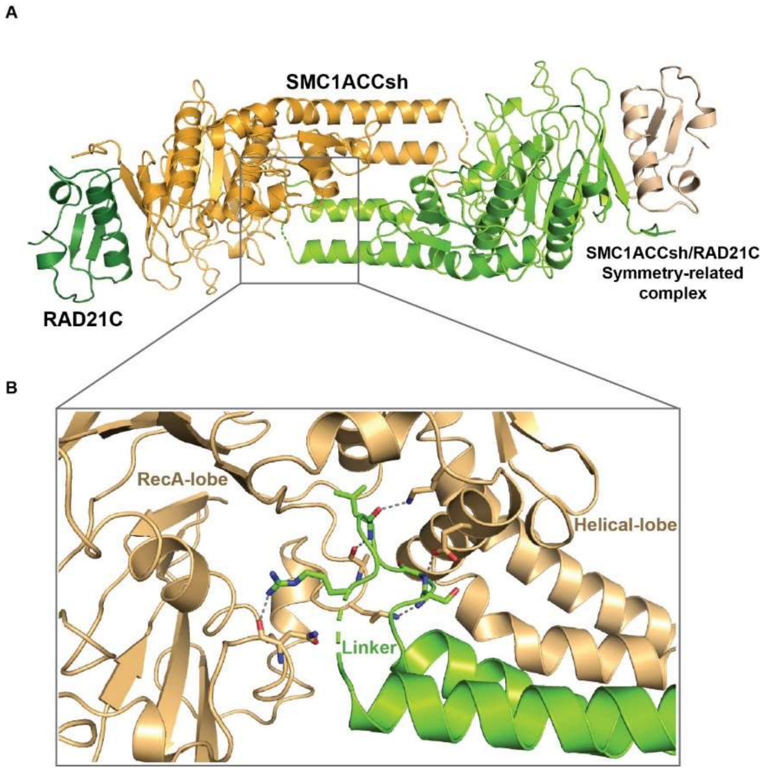


**Supplementary Figure 4. Artificial lever effect induced by the crystal packing of the SMC1ACCsh/RAD21C complex. A.** Crystal packing in the crystals of the SMC1ACCsh/RAD21C complex that occurs regardless of the nucleotide binding state. **B**. Details of the binding of the artificial linker, which bridges the SMC1ACCsh N- and C-terminal coils, at the SMC1ACCsh RecA-lobe/Helical-lobe interface and that induces an artificial lever effect caused by the sliding of the two lobes with respect to each other.

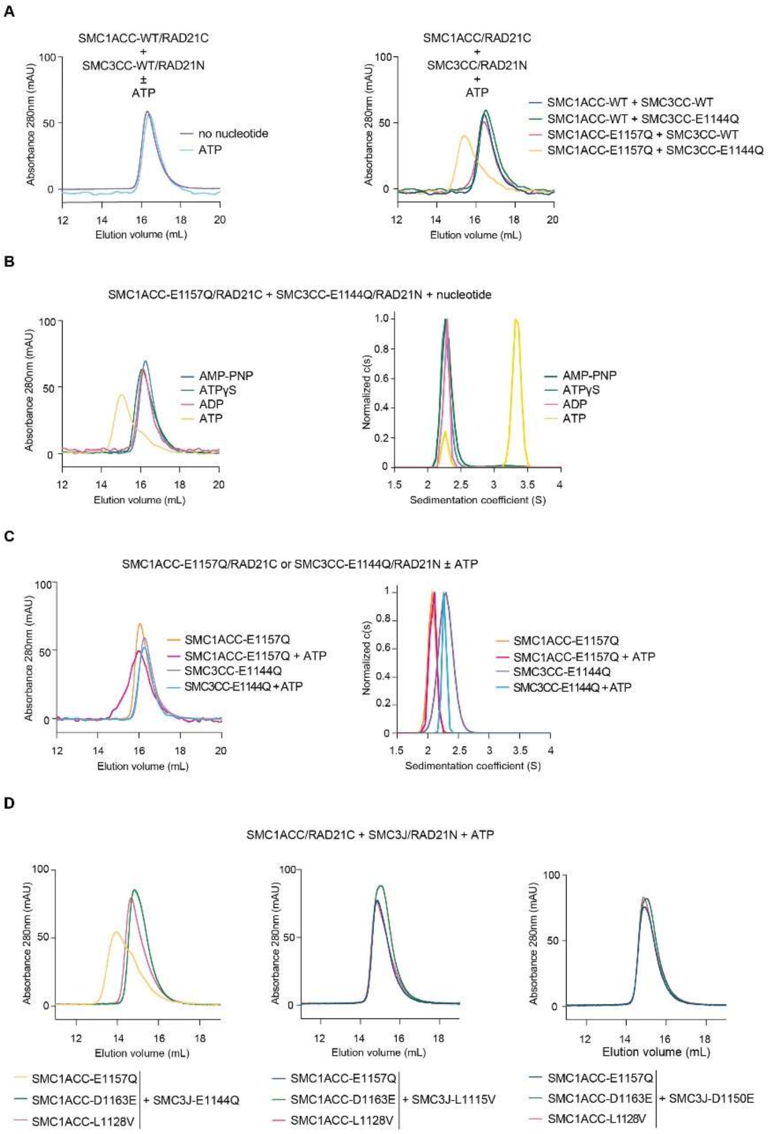


**Supplementary Figure 5. Reconstitution assays of the human ATPase module in the presence of WT and mutant SMC1A and SMC3 HDs and various nucleotides. A.** Size-exclusion chromatography profiles of reconstitution experiments of the human engaged ATPase module using the WT and EQ mutant SMC1ACC/RAD21C and SMC3CC/RAD21N complexes in the absence and presence of ATP. Stable engagement requires both EQ mutants and ATP. **B**. Same as in (A) (left panel) with the EQ mutants and various nucleotides, and confirmation by analytical ultracentrifugation experiments (right panel). Only ATP is able to stably maintain engagement. **C**. Same as in (B) for the characterization of putative SMC1ACC/RAD21C or SMC3CC/RAD21N homodimers in the absence or the presence of ATP. No stable homodimerization is observed. **D**. Same as in (A) using, beside the EQ mutants, the LV and DE mutants that have been used to demonstrate the asymmetry of the Cohesin ATPase active sites. None of the combinations, except that with the two EQ mutants, can stabilize the engaged ATPase module. These experiments have been performed with the SMC3J/RAD21N complex, showing that stable engagement also occurs in the presence of the SMC3 Joint element.

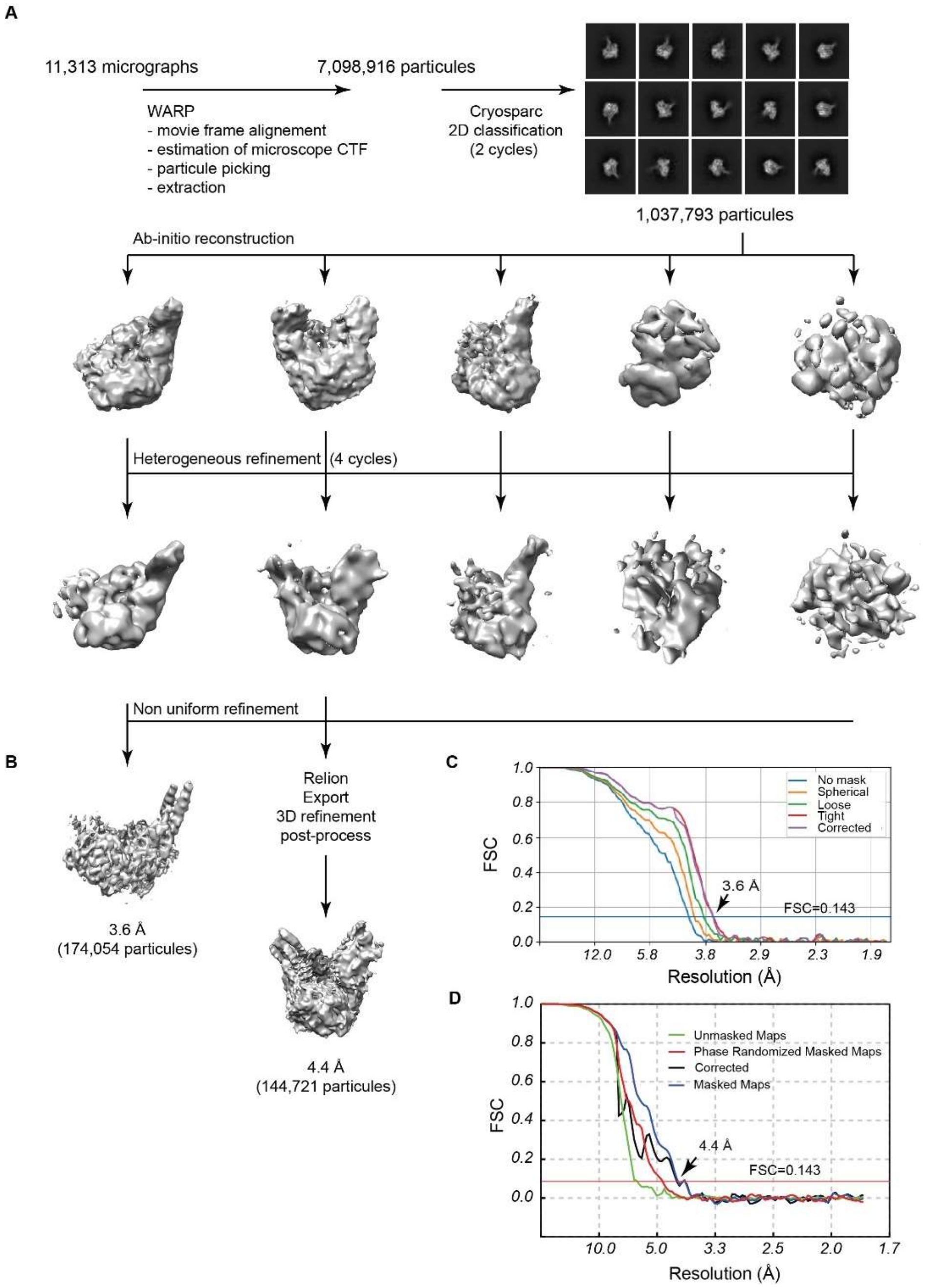


**Supplementary Figure 6. Cryo-EM data processing for the human ATPase module. A.** Flow-chart of the processing of the cryo-EM data obtained on the human engaged ATPase module. The programs and the number of particles used in each step are indicated. The 2D class averages are shown as good classes representatives. **B**. Final cryo-EM maps obtained. The processing led to two usable reconstructions with an average resolution of 3.6 Å and 4.4 Å. The 3.6 Å map was used for initial model building through manual modifications and automated real-space refinement. The model obtained was then used for generating the model at 4.4 Å. **C**. FSC plot for the 3.6 Å cryo-EM map of the human engaged ATPase module. The resolution at which the gold-standard FSC curve drops below the 0.143 threshold is indicated. **D.** Same as in (C) for the 4.4 Å cryo-EM map of the human open-engaged ATPase module.

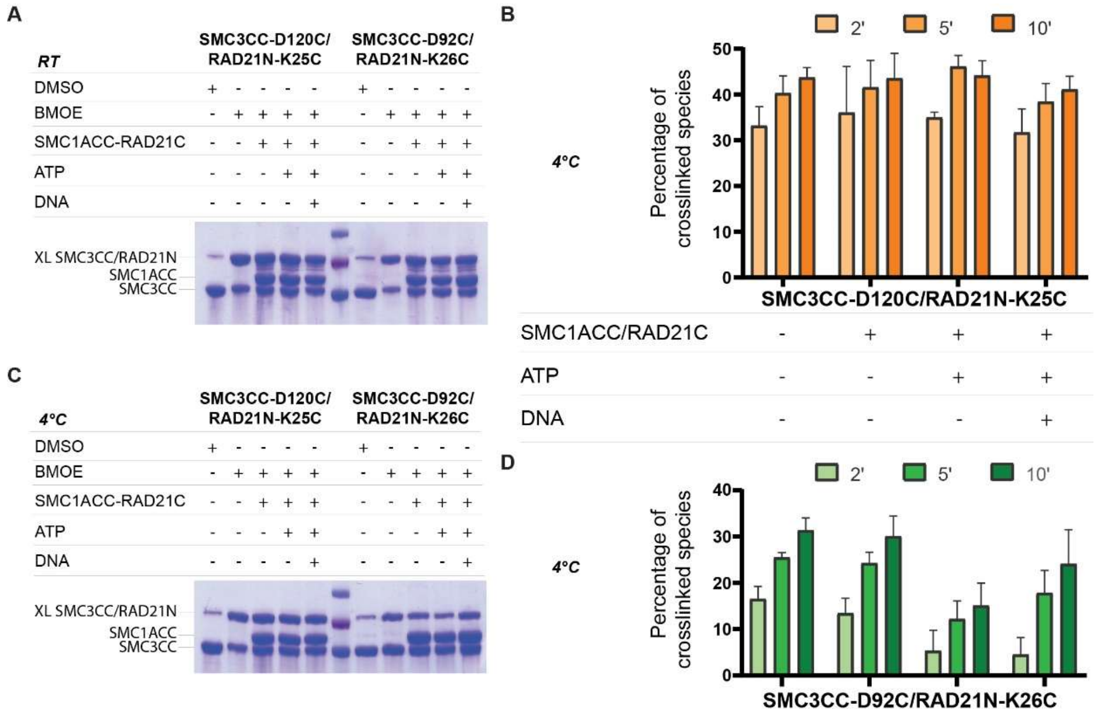


**Supplementary Figure 7. Crosslinking experiments assessing the flexibility of the SMC3 CC/RAD21N complex upon engagement. A.** SDS-PAGE analysis of the 5-minutes crosslinking experiments performed at room temperature (RT) for both crosslinking pairs. XL, crosslinked species. **B**. Quantification of crosslinked species for the SMC3CC-D120C/RAD21N-K25C pair in experiments performed at 4°C. The supplementation of the SMC1ACC/RAD21C, ATP and DNA is indicated underneath the graph. All experiments were done in triplicates. **C**. Same as in (A) with experiments performed at 4°C. **D**. Same as in (C) for the SMC3CC-D92C/RAD21N-K26C pair. The supplementation of the SMC1ACC/RAD21C, ATP and DNA is indicated above the graph in (C). All experiments were done in triplicates.

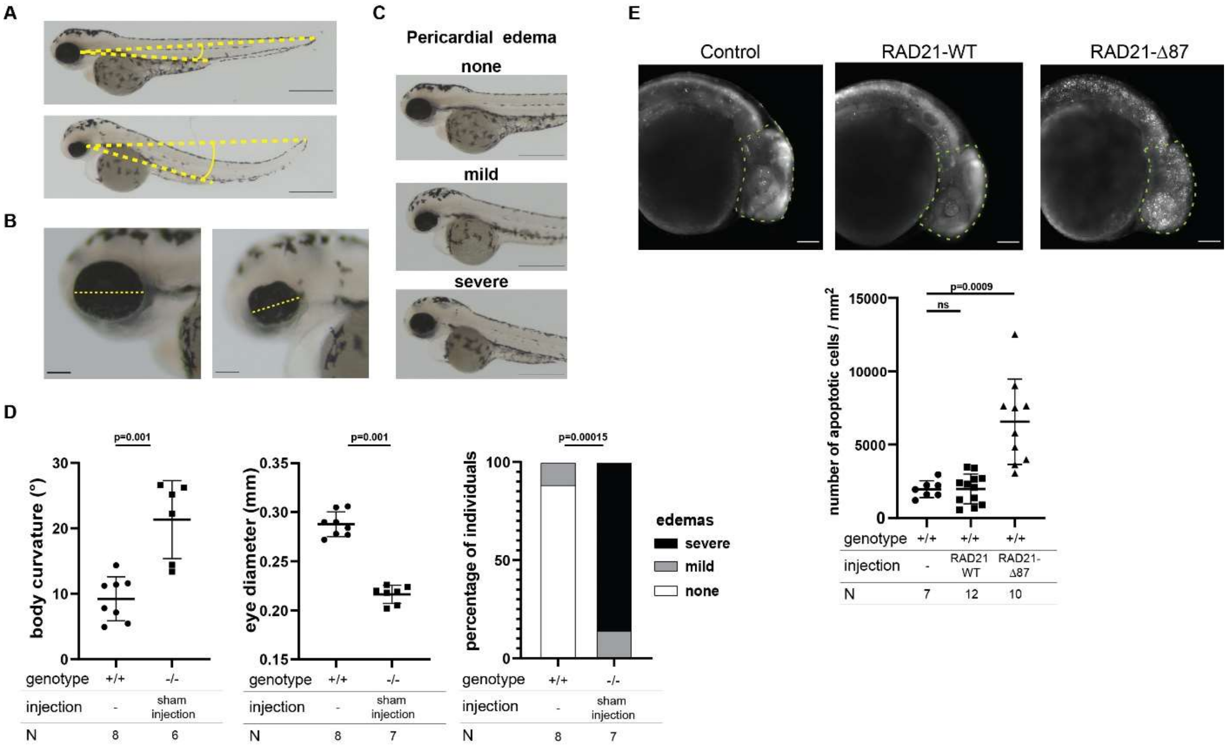


**Supplementary Figure 8. Assessment of the zebrafish body curvature, eye diameter and pericardial edema. A.** Assessment of the body curvature of the zebrafish embryos by measurement of the angle formed between a line going from the tip of the tail to the pigment line that emerges from the lateral side of the eye and a line starting from the same point to the anal disc. Zebrafish larvae with a curved body (lower panel) have a wider angle than WT larvae (upper panel). **B**. Assessment of the zebrafish eye diameter (as assessment of microcephaly) measured from the external side of the eye, where the pigment line emerges, to the opposite side. **C**. Scoring of the zebrafish pericardial edema. From top to bottom, representative images of zebrafish larvae at 3 dpf with no, mild, or severe pericardial edema. **D**. Dot plots showing body curvature (°) and eye diameter (mm) and bar graph showing the presence of pericardial edema for WT and homozygous mutant rad21a$^{hi2529Tg/hi2529Tg}$ larvae. For pericardial edema, larvae were binned into three categories: severe, mild or absent (none) as shown in (C). Statistical significance was assessed by ANOVA followed by a Tukey's test for post-hoc analysis for the body curvature and eye diameter, and a Fisher's exact test for the pericardial edema. All experiments have been performed in duplicates. The p-values are indicated on the graphs. N corresponds to the number of larvae per condition. **E**. From left to right, representative lateral images of zebrafish rad21a+/+ (wildtype; WT) larvae and WT larvae injected with full length WT Danio rerio rad21a (RAD21 WT) or with the rad21a mRNA coding for a mutant lacking the first 87 residues (RAD21-Δ87) at 1-day post-fertilization and stained with acridine orange to visualize necrotic cells. The number of necrotic cells were counted in the highlighted brain area (dotted line on the images). The dot plot represents the number of necrotic cells/mm². A T-test was conducted between pairs of conditions to determine significance. The p-values are indicated on the graphs. Acridine orange staining has been performed two times. N corresponds to the number of embryos per condition. Scale bar, 0.1 mm.

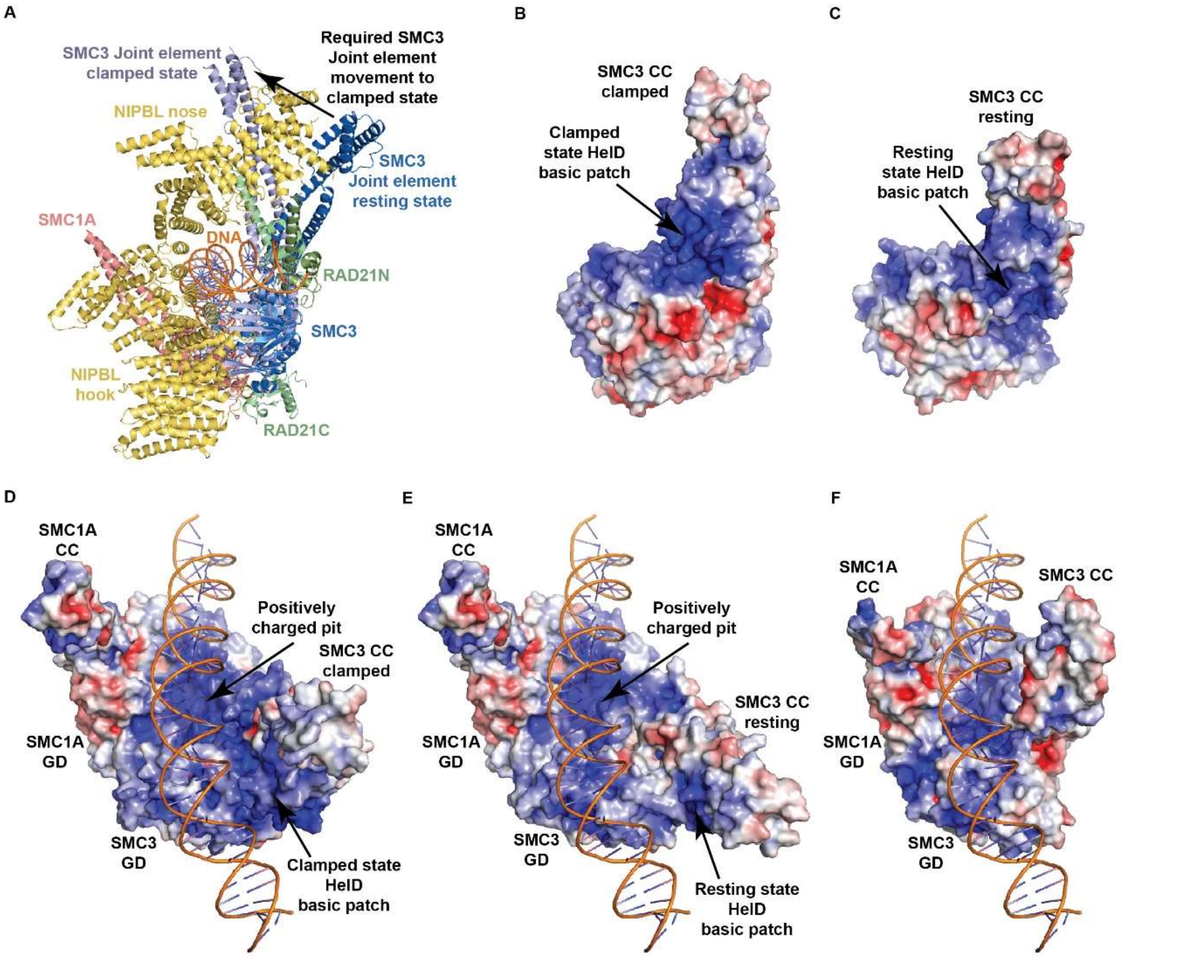


**Supplementary Figure 9. Steric and electrostatic properties of the SMC3C HD/RAD21N complex in the resting and clamped conformations. A.** Ribbon representation of the human clamped complex with the SMC3 CC and Joint element shown both in the clamped and resting conformations (SMC1A HD, pink; SMC3 HD, lavender (clamped)/blue (resting); RAD21, green; NIPBL, yellow; DNA,orange). Only the clamped positioning of the SMC3 Joint is compatible with the positioning of the NIPBL nose in the clamped complex. The resting positioning of the SMC3 Joint as well as any intermediary position clashes with the NIPBL nose, suggesting that both domains have to move conjointly in order to form their clamped interaction. **B.** Electrostatic surface representation of the SMC3CC/RAD21N complex in its clamped conformation (calculations made without the SMC1ACC/RAD21C complex). **C.** Electrostatic surface representation of the resting SMC3CC/RAD21N complex. This stable complex displays a different surface and electrostatic properties than in the clamped conformation shown in (B). **D.** View of DNA bound to the clamped ATPase module represented by its electrostatic surface (red, negatively charged; blue, positively charged). The RAD21N HelD domain is highly positively charged and contributes to the clamping of the DNA in its binding chamber and to its curvature. **E.** Same as in (D) with the SMC3 HD in a resting position. The displacement of the SMC3 CC, and, therefore, of the HelD, widens the DNA binding chamber, potentially facilitating DNA entry and positioning. **F**. Same as in (D) in the case of the open-engaged ATPase module. The conjoint movements of the SMC1A and SMC3 CCs constrict the DNA binding chamber and are not compatible with DNA binding.

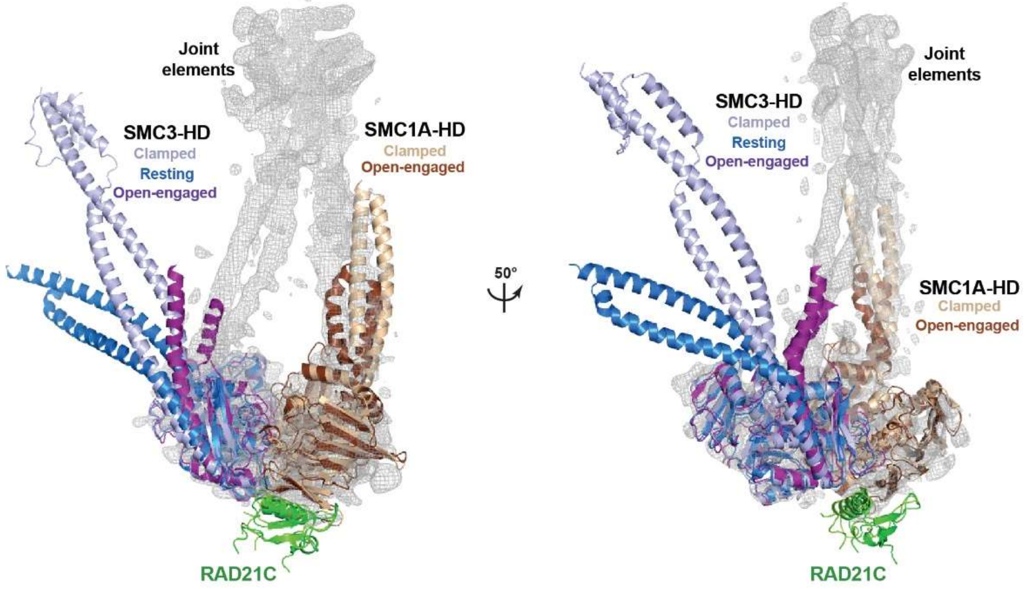


**Supplementary Figure 10. The open-engaged conformation can support an interaction of the SMC1A and SMC3 joint elements while keeping the ATP gate shut.** Fitting of the human resting SMC3 HD, clamped ATPase module and open-engaged ATPase module into the *S. cerevisiae* ATP-engaged Smc1/Smc3 cryo-EM map (EMDB-12889). This modeling indicates that small rearrangements and slight bending of the SMC1A and SMC3 CCs of the open-engaged ATPase module could enable interaction between the SMC1A and SMC3 Joint elements while keeping engagement. Larger movements would be required for the SMC3 CC either in a resting or clamped conformation.

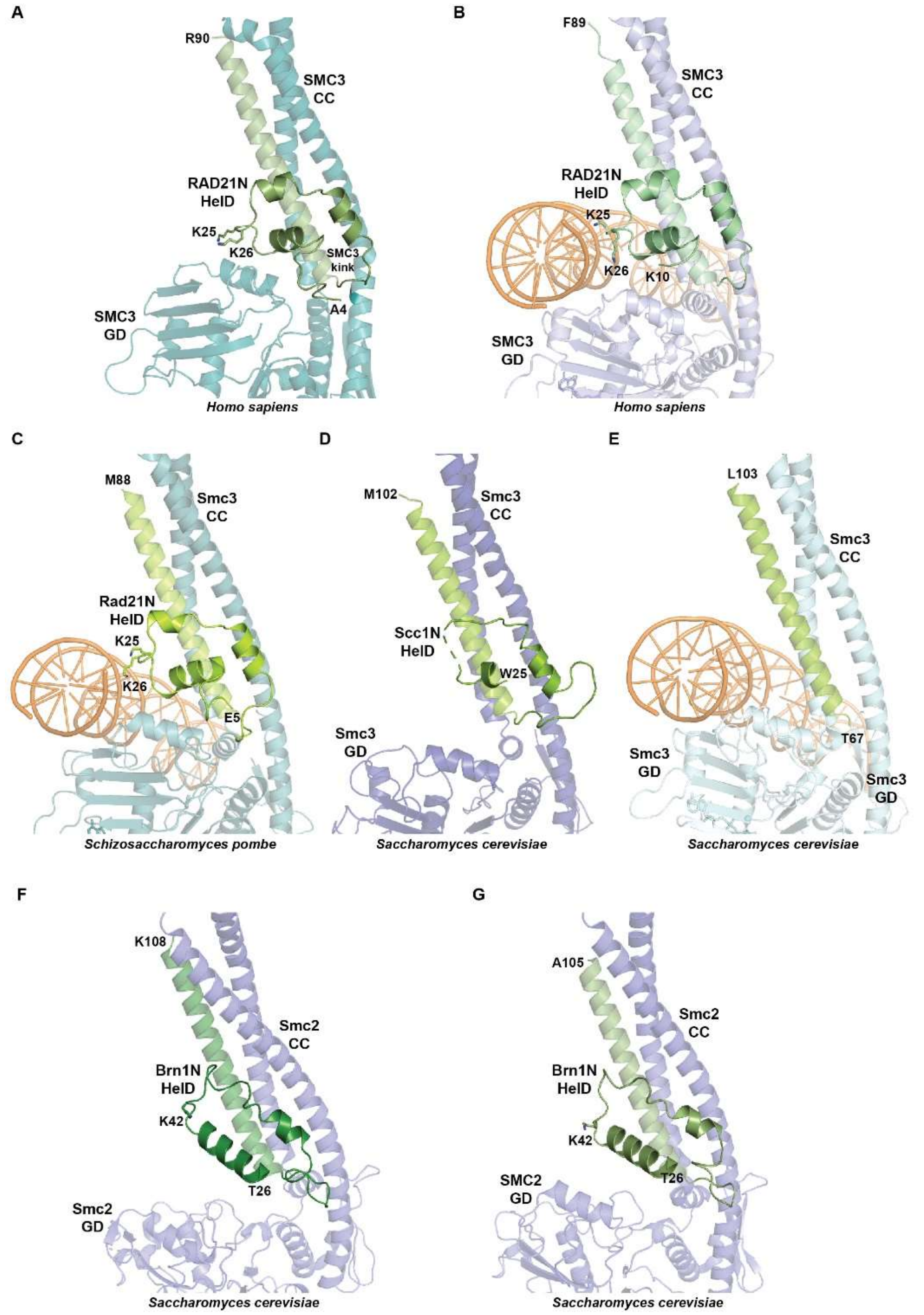


**Supplementary Figure 11. Conformations of the SMC3 HD/RAD21N and Smc2 HD/Brn1N interfaces in Cohesin and Condensin.** Ribbon representations of the SMC3 HD/RAD21N and Smc2 HD/Brn1N interfaces in Cohesin and Condensin. Specific residues, including the N-terminal residue are labeled. DNA is colored orange. **A**. Human Cohesin SMC3 HD/RAD21N complex in the resting conformation (this study). **B**. Human Cohesin SMC3 HD/RAD21N complex in the clamped conformation (PDB entry 6WGE). **C**. *S. pombe* Cohesin Psm3 HD/Rad21N complex in the clamped conformation (PDB entry 6YUF). **D**. *S. cerevisiae* Cohesin Smc3 HD/Scc1N complex in the ATPγS-bound conformation (PDB entry 4UX3). **E**. *S. cerevisiae* Cohesin Smc3 HD/Scc1N complex in the clamped conformation (PDB entry 6ZZ6). **F**. *S. cerevisiae* Condensin Smc2 HD/Brn1N complex in the apo non-engaged conformation (PDB entry 6YVU). **G**. *S. cerevisiae* Condensin Smc2 HD/Brn1N complex in the apo bridged conformation (PDB entry 6YVV).

**Supplementary Table 1. Thermodynamic parameters of ADP, ATP and ATPγS binding to SMC1ACC/RAD21C and SMC3CC/RAD21N measured by Isothermal Titration Calorimetry (ITC)**

| | | | Fitting model: HD + Nucl → HD-Nucl | | | Fitting model: HD + Nucl → HD-Nucl; HD-Nucl + HD-Nucl → HD-Nucl homodimer | | | |
|---|---|---|---|---|---|---|---|---|---|
| Cell | | Syringe | $K_{d1}$ (μM) | $\Delta H_1$ (kcal.mol$^{-1}$) | $N_1$ | $K_{d1}$ (μM) | $\Delta H_1$ (kcal.mol$^{-1}$) | $K_{d2}$ (nM) | $\Delta H_2$ (kcal.mol$^{-1}$) |
| SMC1ACC/RAD21C | wt | ADP | 27.6 ± 7.6 | 2.78 ± 0.19 | 0.85 ± 0.03 | - | - | - | - |
| SMC1ACC/RAD21C | wt | ATP | 51.0 ± 2.58 | 3.17 ± 0.048 | 1.00 ± 0.01 | - | - | - | - |
| SMC1ACC/RAD21C | wt | ATPγS | 31.6 ± 1.00 | 1.89 ± 0.014 | 1.21 ± 0.01 | - | - | - | - |
| SMC1ACC/RAD21C | E1157Q | ATP | 55.3 ± 4.72 | 1.42 ± 0.038 | 1.05 ± 0.01 | - | - | - | - |
| SMC1ACC/RAD21C | E1157Q | ATPγS | 43.0 ± 3.06 | 1.06 ± 0.021 | 1.01 ± 0.01 | - | - | - | - |
| SMC3CC/RAD21N | wt | ADP | 1.8 ± 0.07 | -7.19 ± 0.02 | 0.98 ± 0.02 | - | - | - | - |
| SMC3CC/RAD21N | wt | ATP | - | - | - | 51.1 ± 23.7 | -2.7 ± 1.2 | 1.4 ± 0.7 | -18 ± 0.8 |
| SMC3CC/RAD21N | wt | ATPγS | - | - | - | 18.3 ± 1.5 | -6.0 ± 1.9 | 5.3 ± 0.6 | -7.4 ± 1.8 |
| SMC3CC/RAD21N | E1144Q | ATP | - | - | - | 74.6 ± 33.8 | -5.2 ± 1.0 | 2.8 ± 1.8 | -14.9 ± 0.6 |
| SMC3CC/RAD21N | E1144Q | ATPγS | - | - | - | 35.5 ± 9.2 | -5.6 ± 0.8 | 3.8 ± 1.0 | -8.0 ± 0.7 |

**Supplementary Table 2. Crystallization conditions and crystallographic statistics for the SMC1ACC/RAD21C complexes.**

| SMC1ACC/ RAD21C | WT apo | EQ mutant apo | WT loop |
|---|---|---|---|
| **Crystallization conditions** | 0.1 M Hepes pH 7.0; 10% w/v PEG 6000 | 0.2 M sodium malonate dibasic monohydrate; 0.1 M Bis-Tris propane pH 6.5; 20% w/v PEG 3350 | 0.1 M Hepes pH 7.0; 10% w/v PEG 6000 |
| **Data collection*** | | | |
| Space group | C2 | C2 | C2 |
| Cell dimensions | | | |
| a, b, c (Å) | 187.93, 64.72, 47.34 | 158.55, 67.07, 51.63 | 187.60, 64.59, 47.13 |
| α, β, γ (°) | 90.00, 103.07, 90.00 | 90.00, 92.32, 90.00 | 90.00, 103.97, 90.00 |
| Resolution (Å) | 50.00 – 2.20 (2.34 – 2.20) | 50.00 – 1.90 (2.02 – 1.90) | 50.00 – 2.09 (2.22 – 2.09) |
| Rsym or Rmerge | 0.072 (1.196) | 0.091 (1.545) | 0.06 (1.206) |
| I / σI | 12.08 (1.05) | 9.56 (1.02) | 12.64 (1.32) |
| Completeness (%) | 98.6 (87.3) | 99.7 (98.3) | 98.9 (95.5) |
| Redundancy | 4.9 (4.4) | 6.8 (6.9) | 4.6 (4.6) |
| CC(1/2) | 99.8 (51.3) | 99.8 (71.9) | 99.9 (72.0) |
| **Refinement** | | | |
| Resolution (Å) | 46.12 – 2.20 | 44.05 – 1.90 | 32.30 – 2.09 |
| No. reflections | 27871 | 42461 | 32074 |
| Rwork / Rfree | 0.190 / 0.244 | 0.195 / 0.231 | 0.199 / 0.223 |
| Number of atoms | | | |
| Protein | 3248 | 3288 | 3136 |
| Ligand/ion | - | - | - |
| Water | 79 | 132 | 52 |
| B-factors ($Å^2$) | | | |
| Protein | 72.77 | 62.17 | 82.58 |
| Ligand/ion | - | - | - |
| Water | 58.40 | 53.06 | 58.55 |
| R.m.s. deviations | | | |
| Bond lengths (Å) | 0.008 | 0.006 | 0.008 |
| Bond angles (º) | 0.913 | 0.771 | 0.905 |

* *Values in parentheses are for the highest-resolution shell.*

| SMC1ACC/ RAD21C | EQ mutant Loop | EQ mutant ADP | EQ mutant ATPγS-Mg |
|---|---|---|---|
| **Crystallization conditions** | 0.2 M sodium bromide; 0.1 M Bis-Tris propane pH 6.5; 20% w/v PEG 3350 | 0.2 M ammonium formate; 20% w/v PEG 3350 | 0.1 M MMT pH 7.0; 25% w/v PEG 1500 |
| **Data collection*** | | | |
| Space group | C2 | C2 | $P4_1 2_1 2$ |
| Cell dimensions | | | |
| a, b, c (Å) | 187.92, 64.60, 89.95 | 189.31, 65.28, 105.76 | 68.92, 68.92, 215.33 |
| α, β, γ (°) | 90.00, 99.74, 90.00 | 90.00, 116.66, 90.00 | 90.00, 90.00, 90.00 |
| Resolution (Å) | 50.00 – 1.77 (1.87 – 1.77) | 50.00 – 2.44 (2.58 – 2.44) | 50.00 – 2.50 (2.65 – 2.50) |
| Rsym or Rmerge | 0.055 (1.277) | 0.120 (1.121) | 0.126 (3.602) |
| I / σI | 15.94 (1.05) | 11.60 (1.26) | 19.14 (0.84) |
| Completeness (%) | 99.7 (98.7) | 99.5 (98.3) | 98.4 (98.5) |
| Redundancy | 6.8 (6.2) | 6.4 (5.2) | 24.7 (24.9) |
| CC(1/2) | 99.9 (58.0) | 99.8 (57.9) | 100.0 (53.8) |
| **Refinement** | | | |
| Resolution (Å) | 44.63 – 1.77 | 47.80 – 2.44 | 48.74 – 2.50 |
| No. reflections | 103917 | 43312 | 18516 |
| Rwork / Rfree | 0.190 / 0.218 | 0.194 / 0.250 | 0.221 / 0.298 |
| Number of atoms | | | |
| Protein | 6469 | 6578 | 3312 |
| Ligand/ion | 19 | 54 | 32 |
| Water | 389 | 64 | 32 |
| B-factors ($Å^2$) | | | |
| Protein | 47.11 | 60.76 | 93.06 |
| Ligand/ion | 62.46 | 57.85 | 67.92 |
| Water | 46.96 | 51.23 | 81.54 |
| R.m.s. deviations | | | |
| Bond lengths (Å) | 0.007 | 0.008 | 0.008 |
| Bond angles (º) | 0.886 | 0.965 | 1.024 |
| ** Values in parentheses are for the highest-resolution shell.* | | | |

**Supplementary Table 3. Crystallization conditions and crystallographic statistics for the SMC1ACCsh/RAD21C complexes.**

| SMC1ACCsh/ RAD21C | EQ mutant Apo | EQ mutant Loop | EQ mutant ADP |
|---|---|---|---|
| **Crystallization conditions** | 0.2 M ammonium nitrate pH 6.3; 20% w/v PEG 3350 | 0.2 M sodium sulfate; 20 % w/v PEG 3350 | 0.2 M sodium malonate; 0.1 M Bis Tris propane pH 6.5; 20 % w/v PEG 3350 |
| **Data collection*** | | | |
| Space group | I 2 2 2 | I 2 2 2 | I 2 2 2 |
| Cell dimensions | | | |
| a, b, c (Å) | 66.14, 114.04, 134.87 | 66.11, 113.84, 134.47 | 66.04, 113.84, 134.19 |
| α, β, γ (°) | 90.00, 90.00, 90.00 | 90.00, 90.00, 90.00 | 90.00, 90.00, 90.00 |
| Resolution (Å) | 50.00 - 1.85 (1.96 – 1.85) | 50.00 – 1.50 (1.59 – 1.50) | 50.00 – 1.36 (1.44 -1.36) |
| Rsym or Rmerge | 0.078 (1.308) | 0.087 (2.382) | 0.067 (2.855) |
| I / σI | 21.22 (1.68) | 17.87 (1.08) | 20.31 (0.92) |
| Completeness (%) | 99.9 (99.2) | 99.9 (99.4) | 99.9 (99.7) |
| Redundancy | 12.7 (10.1) | 13.2 (12.8) | 13.2 (13.5) |
| CC(1/2) | 100.0 (75.6) | 100.0 (43.2) | 100.0 (38.7) |
| **Refinement** | | | |
| Resolution (Å) | 43.63 – 1.85 | 43.55 – 1.50 | 43.49 – 1.36 |
| No. reflections | 43937 | 81150 | 108065 |
| Rwork / Rfree | 0.181 / 0.222 | 0.198 / 0.232 | 0.204 / 0.219 |
| Number of atoms | | | |
| Protein | 3397 | 3313 | 3413 |
| Ligand/ion | - | 5 | 27 |
| Water | 257 | 360 | 301 |
| B-factors ($Å^2$) | | | |
| Protein | 41.05 | 32.12 | 31.19 |
| Ligand/ion | - | 22.86 | 47.72 |
| Water | 46.15 | 42.17 | 38.25 |
| R.m.s. deviations | | | |
| Bond lengths (Å) | 0.006 | 0.006 | 0.005 |
| Bond angles (º) | 0.809 | 0.831 | 0.833 |

** Values in parentheses are for the highest-resolution shell.*

| SMC1ACCsh/ RAD21C | EQ mutant ADP-Mg | EQ mutant AGS-Mg |
|---|---|---|
| **Crystallization conditions** | 0.2 M magnesium chloride; 0.1 M MES pH 6.0; 20 % w/v PEG 6000 | 0.2 M sodium formate; 0.1 M Bis Tris propane pH 6.5; 20 % w/v PEG 3350 |
| **Data collection*** | | |
| Space group | I 2 2 2 | I 2 2 2 |
| Cell dimensions | | |
| a, b, c (Å) | 66.45, 113.91, 133.84 | 66.22, 113.98, 134.07 |
| α, β, γ (°) | 90.00, 90.00, 90.00 | 90.00, 90.00, 90.00 |
| Resolution (Å) | 50.00 – 1.65 (1.75 – 1.65) | 50.00 – 1.94 (2.06– 1.94) |
| Rsym or Rmerge | 0.113 (2.391) | 0.166 (2.378) |
| I / σI | 15.52 (1.05) | 10.95 (1.06) |
| Completeness (%) | 99.8 (99.1) | 99.8 (99.0) |
| Redundancy | 13.3 (13.2) | 12.9 (11.7) |
| CC(1/2) | 99.9 (42.0) | 99.9 (36.6) |
| **Refinement** | | |
| Resolution (Å) | 43.57 - 1.65 | 43.54 - 1.94 |
| No. reflections | 61052 | 37968 |
| Rwork / Rfree | 0.196 / 0.228 | 0.205 / 0.253 |
| Number of atoms | | |
| Protein | 3401 | 3289 |
| Ligand/ion | 28 | 32 |
| Water | 302 | 212 |
| B-factors (Å$^2$) | | |
| Protein | 33.63 | 43.99 |
| Ligand/ion | 47.22 | 64.80 |
| Water | 42.81 | 46.12 |
| R.m.s. deviations | | |
| Bond lengths (Å) | 0.007 | 0.007 |
| Bond angles (º) | 0.829 | 0.881 |

* *Values in parentheses are for the highest-resolution shell.*

**Supplementary Table 4. Crystallization conditions and crystallographic statistics for the SMC3CC/RAD21N complexes.**

| SMC3CC/ RAD21N | EQ mutant Apo | EQ mutant ADP | WT ADP-Mg |
|---|---|---|---|
| **Crystallization conditions** | 0.21M $NaSO_4$; 0.1M Bis-tris propane pH7; 16% w/v PEG3350 | 0.1 M Na/K phosphate pH 7.5; 0.1 M HEPES pH 7.5; 15% v/v PEG Smear High; 10% v/v ethylene glycol | 0.075 M magnesium chloride; 0.075 M Sodium citrate tribasic dehydrate; 0.1 M Bis-Tris pH 6.0; 18% v/v PEG Smear Broad |
| **Data collection*** | | | |
| Space group | $P4_1 2_1 2$ | $P4_1 2_1 2$ | $P4_1 2_1 2$ |
| Cell dimensions | | | |
| a, b, c (Å) | 90.79, 90.79, 236.87 | 89.95, 89.95, 233.91 | 89.71, 89.71, 234.65 |
| α, β, γ (°) | 90.00, 90.00, 90.000 | 90.00, 90.00, 90.000 | 90.00, 90.00, 90.000 |
| Resolution (Å) | 50.00 – 2.60 (2.76 – 2.60) | 50.00 – 2.45 (2.60 – 2.45) | 50.00 – 3.00 (3.18 – 3.00) |
| Rsym or Rmerge | 0.280 (3.842) | 0.154 (2.621) | 0.247 (3.684) |
| I / σI | 14.93 (1.02) | 22.60 (1.50) | 16.53 (1.11) |
| Completeness (%) | 99.9 (99.7) | 99.9 (99.7) | 99.9 (99.6) |
| Redundancy | 26.2 (25.7) | 26.1 (26.4) | 26.0 (26.9) |
| CC(1/2) | 99.9 (36.6) | 99.9 (58.1) | 99.9 (43.1) |
| **Refinement** | | | |
| Resolution (Å) | 49.60 – 2.60 | 49.03 - 2.45 | 49.27 – 3.00 |
| No. reflections | 31408 | 36209 | 19996 |
| Rwork / Rfree | 0.215 / 0.252 | 0.195 / 0.239 | 0.196 / 0.264 |
| Number of atoms | | | |
| Protein | 4040 | 3918 | 3931 |
| Ligand/ion | 11 | 27 | 28 |
| Water | 9 | 110 | 4 |
| B-factors ($Å^2$) | | | |
| Protein | 71.43 | 74.80 | 96.65 |
| Ligand/ion | 75.57 | 55.69 | 79.86 |
| Water | 53.35 | 62.44 | 71.39 |
| R.m.s. deviations | | | |
| Bond lengths (Å) | 0.008 | 0.008 | 0.009 |
| Bond angles (º) | 0.906 | 0.940 | 1.080 |

** Values in parentheses are for the highest-resolution shell.*

| SMC3CC/ RAD21N | EQ mutant ATPγS-Mg (form 1) | EQ mutant ATPγS-Mg (form 2) |
|---|---|---|
| Crystallization conditions | 0.3 M NaCl; 0.05 M L-Arginine; 0.1 M Tris pH7.5; 22.5 % v/v PEG Smear Broad; 0.05 M L-Glutamic acid monosodium salt hydrate | 0.1 M Sodium chloride; 0.1 M Bicine pH 9.0; 20% w/v PEG 550 MME |
| **Data collection*** | | |
| Space group | $C2\ 2\ 2_1$ | $P6_4\ 2\ 2$ |
| Cell dimensions | | |
| a, b, c (Å) | 118.24, 154.46, 136.54 | 201.07, 201.07, 92.38 |
| α, β, γ (°) | 90.00, 90.00, 90.00 | 90.00, 90.00, 90.00 |
| Resolution (Å) | 50.00 – 2.25 (2.39 - 2.25) | 50.00 – 3.11 (3.30 – 3.11) |
| Rsym or Rmerge | 0.098 (2.233) | 0.256 (4.814) |
| I / σI | 20.32 (1.34) | 17.00 (1.03) |
| Completeness (%) | 98.0 (94.2) | 99.5 (98.1) |
| Redundancy | 13.6 (13.1) | 39.1 (39.1) |
| CC(1/2) | 100.0 (81.6) | 99.9 (63.7) |
| **Refinement** | | |
| Resolution (Å) | 47.21 – 2.25 | 44.64 – 3.11 |
| No. reflections | 58103 | 20146 |
| Rwork / Rfree | 0.202 / 0.253 | 0.187 / 0.238 |
| Number of atoms | | |
| Protein | 7529 | 3955 |
| Ligand/ion | 64 | 32 |
| Water | 90 | - |
| B-factors (Å$^2$) | | |
| Protein | 75.14 | 112.95 |
| Ligand/ion | 47.68 | 93.80 |
| Water | 56.96 | - |
| R.m.s. deviations | | |
| Bond lengths (Å) | 0.008 | 0.008 |
| Bond angles (º) | 0.927 | 0.997 |

** Values in parentheses are for the highest-resolution shell.*

**Supplementary Table 5. Cryo-EM data collection, refinement and validation statistics for the engaged and open-engaged ATPase modules.**

| **Data collection and processing** | **Collection 1 (PEGylated gold 1.2/1.3 grid)** | **Collection 2 (Gold 1.2/1.3 grid)** | **Collection 3 (C-flat 1.2/1.3)** |
|---|---|---|---|
| Microscope and Camera | Glacios$^{TM}$ cryo-TEM, K2 Summit | | |
| Magnification | 45,000x | | |
| Voltage (kV) | 200 | | |
| Number of micrographs | 4146 | 1593 | 5574 |
| Electron exposure (e$^-$/Å$^2$) | 44.42 | 43.61 | 63.85 |
| Defocus range | -0.8 to -2.0 | -0.8 to -2.0 | -0.8 to -3.0 |
| Pixel size | 0.901 | | |
| Symmetry imposed | C1 | | |
| Number of initial particles | 2,367,926 | 2,119,775 | 2,611,215 |

| **Processing** | **ATPase module engaged** | **ATPase module open engaged** |
|---|---|---|
| Number of final particles | 174,054 | 144,721 |
| Map resolution (Å) | 3.6 | 4.4 |
| FSC threshold | 0.143 | 0.143 |
| **Refinement** | | |
| Model resolution range (Å) | 3.6 | 4.4 |
| FSC threshold | 0.143 | 0.143 |
| Map sharpening *B* factor (Å$^2$) | 182.6 | -164.7 |
| *Model composition* | | |
| Non-H atoms | 5764 | 6300 |
| Protein residues | 718 | 782 |
| Ligands | 4 | 2 |
| B factors (Å$^2$) | | |
| Protein | 99.81 | 305.62 |
| Ligand | 73.49 | 230.23 |
| *r.m.s. deviations* | | |
| Bond lengths | 0.003 | 0.003 |
| Bond angles | 0.641 | 0.664 |
| **Validation** | | |
| MolProbity score | 2.17 | 2.42 |
| Clashscore | 19.37 | 25.10 |
| Poor rotamers (%) | 0 | 0 |
| *Ramachandran plot* | | |
| Favored (%) | 94.33 | 90.76 |
| Allowed (%) | 5.67 | 9.24 |
| Outlier (%) | 0.0 | 0.0 |
